# The Ca^2+^ sensor protein CMI1 fine tunes root development, auxin distribution and responses

**DOI:** 10.1101/488536

**Authors:** Ora Hazak, Elad Mamon, Meirav Lavy, Hasana Sternberg, Smrutisanjita Behera, Ina Schmitz-Thom, Daria Bloch, Olga Dementiev, Itay Gutman, Tomer Danziger, Netanel Schwarz, Anas Abuzeineh, Keithanne Mockaitis, Mark Estelle, Joel A. Hirsch, Jörg Kudla, Shaul Yalovsky

## Abstract

Signaling cross-talks between auxin, a regulator of plant development and Ca^2+^, a universal second messenger have been proposed to modulate developmental plasticity in plants. However, the underlying molecular mechanisms are largely unknown. Here we report that in *Arabidopsis* roots, auxin elicits specific Ca^2+^ signaling pattern that spatially coincide with the expression pattern of auxin-regulated genes. We identified the EF-hand protein CMI1 (Ca^2+^ sensor Modulator of ICR1) as an interactor of the ROP effector ICR1 (Interactor of Constitutively active ROP). CMI1 is monomeric in solution, changes its secondary structure at Ca^2+^ concentrations ranging from 10^−9^ to 10^−8^ M and its interaction with ICR1 is Ca^2+^ dependent, involving a conserved hydrophobic pocket. *cmi1* mutants display an increased auxin response including shorter primary roots, longer root hairs, longer hypocotyls and altered lateral root formation while ectopic expression of CMI1 induces root growth arrest and reduced auxin responses at the root tip. When expressed alone, CMI1 is localized at the plasma membrane, the cytoplasm and in nuclei. Interaction of CMI1 and ICR1 results in exclusion of CMI1 from nuclei and suppression of the root growth arrest. CMI1 expression is directly upregulated by auxin while expression of auxin induced genes is enhanced in *cmi1* concomitantly with repression of auxin induced Ca^2+^ increases in the lateral root cap and vasculature, indicating that CMI1 represses early auxin responses. Collectively, our findings identify a crucial function of Ca^2+^ signaling and CMI1 in root growth and suggest an auxin-Ca^2+^ regulatory feedback loop that fine tunes root development.

## Introduction

The plant hormone auxin functions as a morphogen by forming local maxima and gradients and regulates diverse developmental and physiological processes (1). Auxin operates as a “molecular glue” mediating the binding of the Aux/IAA transcriptional repressors to the Skp Cullin F-box Transport Inhibitor Response 1 (SCF^TIR1/AFB^) E3 ubiquitin ligase complex, resulting in polyubiquitilation and proteasomal degradation of the Aux/IAAs, leading to activation of the ARF (Auxin Response Factor) transcriptional regulators (1–4). In addition, auxin induces rapid transcription-independent responses such as membrane depolarization and Ca^2+^ influx by mechanisms that depend on auxin perception by TIR1/AFB (5–9). Signaling cross-talks between auxin and Ca^2+^have been proposed to modulate developmental plasticity in plants (5, 10–12). Ca^2+^ is a universal second messenger that transduces exogenous and endogenous signals to trigger cellular and developmental responses (13, 14). Considering its diverse effects, Ca^2+^ has been named “the missing link in auxin action” (15). AUX1-dependent auxin influx in root and root hairs induces CNGC14- and TIR1/AFB-dependent Ca^2+^ signaling within seconds that in turn affects downstream auxin signaling (5–7). Cyclic Nucleotide Gated Channel 14 (CNGC14) function is required in response to gravity stimulus (6), indicating that function involves Ca^2+^ signaling.

Auxin transport depends on AUX1/LAX auxin influx transporters (16), PINFORMED (PIN) proteins, ABCB auxin efflux transporters (17, 18) and under low nitrogen conditions by NRT1.1 NO_3_^−^ influx transporter (19, 20). The AGCVIII kinase PINOID (PID), which regulates PIN1, PIN2 and PIN3 distribution (21, 22) and PIN mediated auxin transport (23), interacts with two EF-hand Ca^2+^ binding proteins, TOUCH3 (TCH3) and PID Binding Protein 1 (PBP1) (24). Moreover, PID overexpression-induced root meristem collapse was reduced by treatments with LaCl_3_, a Ca^2+^ channel inhibitor suggesting the requirement of Ca^2+^ for PID function and consequently PIN regulation (24). However, it is not known yet how the Ca^2+^ binding proteins TCH3 and PBP1 affect PID function.

The role of Ca^2+^ signaling and Ca^2+^ binding protein(s) that transduce auxin-related Ca^2+^ signals is only partially understood. Therefore, the mechanistic basis for the interplay of auxin and Ca^2+^ signaling is not well known. In this work we describe the identification of an auxin regulated Ca^2+^ binding protein that crucially regulates auxin responses and affects auxin-induced changes in cytoplasmic Ca^2+^ levels.

## Results

### Auxin induces specific Ca^2+^ signal pattern in the root

To study potential changes in cytoplasmic Ca^2+^ concentration in the root following auxin treatment, we used Arabidopsis seedlings expressing the FRET-based Ca^2+^ indicator Yellow Cameleon 3.6 (YC3.6) (25). Time-lapse imaging was performed in 5-7d old *Arabidopsis* roots by exchanging control buffer to buffer containing 10 μM naphthaleneacetic acid (NAA) using epifluorescent (Fig 1A) or confocal microscopes (Fig 1B and C). In control conditions, elevated Ca^2+^ concentrations were primarily observed in the QC, the proximal layer of the columella, the lateral root cap (LRC) and vascular tissues (Fig 1A (mock) and 1B (before treatment, overview). A typical auxin-induced Ca^2+^ signal was observed after one minute of auxin (10 μM NAA) application. The most pronounced Ca^2+^ elevations were observed in the root cap, lateral root cap and vasculature (Fig 1A-C, NAA). The pattern of the generated Ca^2+^ signal was corresponding to auxin response and distribution (26–28). The similarity between auxin induced Ca^2+^ concentration increases (Fig 1A) and the oscillatory expression pattern of TIR1/AFB auxin receptors regulated genes (26) is suggestive for mutual interdependency between auxin and Ca^2+^ in the root.

**Fig 1.**
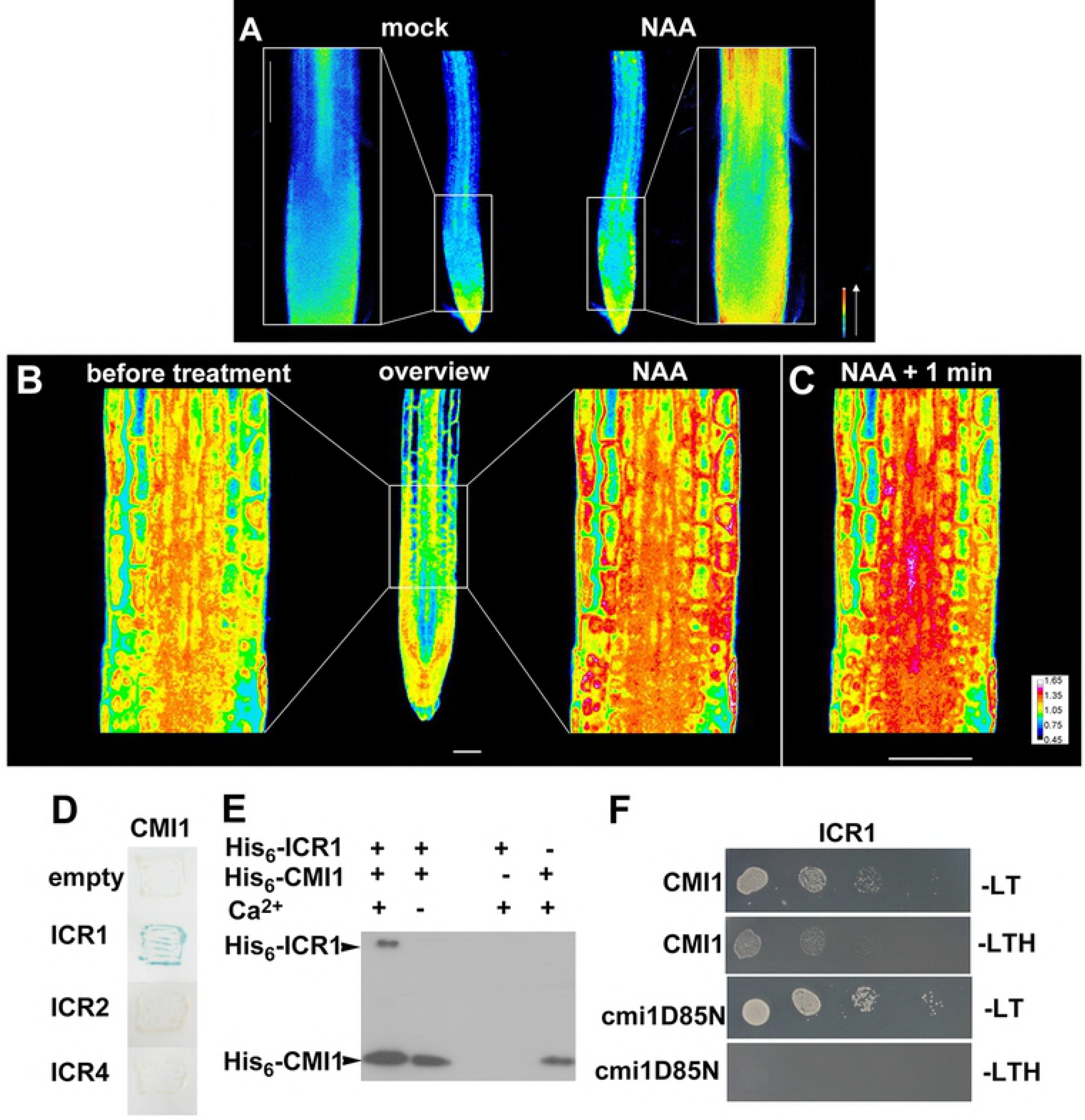
The auxin regulated Ca^2+^-binding protein CMI1 interacts with ICR1 in a Ca^2+^ dependent fashion. (A) Epifluorecscent images of root expressing the Yellow Cameleon (YC3.6) free Ca^2+^ sensor prior to auxin treatment (mock) and 100 seconds after treatment with 10 μM NAA (NAA). (B) Confocal images of root expressing the Yellow Cameleon (YC3.6) free Ca^2+^ sensor prior to auxin treatment (before treatment) and 100 seconds after treatment with 10 μM NAA (NAA) (C) The same root shown in B imaged after additional 60 second (NAA + 1 min). (D) CMI1 interacts with ICR1 but not with ICR2 or ICR4 in yeast two-hybrid assays. (E) Protein immuno blot decorated with anti polyHis-tag monoclonal antibodies showing that co-immunoprecipitation of His-CMI1 and His-ICR1 is Ca^2+^ dependent. (F) ICR1 interacts with CMI1 but not with the cmi1D85N Ca^2+^ non-binding mutant in yeast two-hybrid assays. -LT: Leu, Trp deficient medium; -LTH: Leu, Trp, His deficient medium. Scale bars, 20 μm.

Previously, we identified a family of coiled coil domain ROP (Rho Of Plants) effectors that we named ICRs (Interactor of Constitutively active ROP) (29). ICR1 regulates cell polarity, is degraded in an auxin dependent fashion in the root meristem an affects root growth (29–31). In a screen for ICR1 interacting proteins we identified a single EF-hand Ca^2+^ binding protein that we designated as CMI1 (Ca^2+^ sensor Modulator of ICR1) (*At4g27280*). CMI1 is a small 14 kDa protein containing a single EF-hand (S1A Fig). In *Arabidopsis* CMI1 is a member of a small protein family consisting of 3 members and was formerly called KRP1 (KIC Related Protein 1) (32). Because the name KRP has originally been used for the cell cycle regulators KIP Related Proteins (33), which are unrelated to KRP1, we decided to adhere to the CMI1 nomenclature in this work.

### Ca^2+^ promotes the interaction between CMI1 and ICR1

CMI1 interacted specifically with ICR1 but not with ICR2 (*At2g37080*) or ICR4 (*At1g78430*), the closest homologues of ICR1 (Fig 1D). To further characterize the interaction between CMI1 and ICR1 and to examine whether it is Ca^2+^-dependent, we performed *in vitro* pull-down experiments. His-ICR1 was immunoprecipitated together with His-CMI1 using anti CMI1 antibodies in the presence of Ca^2+^ (Fig 1E). In contrast, pull down of His-CMI1 by GST-ICR1 did not take place when the Ca^2+^ was chelated with EGTA. The interaction of ICR1 and CMI1 in the pull-down assays was specific since His-CMI1 was not precipitated by non-fused GST or glutathione beads (S1B and C Fig). To further corroborate that the interaction between ICR1 and CMI1 is Ca^2+^ dependent, we created a CMI1 D85N mutant in which a conserved EF-hand Asp required for Ca^2+^ binding (34) was mutated to Asn (S1A Fig). Yeast two-hybrid assays showed that CMI1 interacts with ICR1 but not with the CMI1D85N protein (Fig 1F). Taken together, these results establish that the interaction between ICR1 and CMI1 is Ca^2+^ dependent both in yeast and *in vitro*.

### CMI1 functions as a monomeric Ca^2+^ sensor

Circular dichroism spectroscopy (CD-spec) was used to examine changes in CMI1 secondary structure at different free Ca^2+^ concentrations. The analysis was carried out in solutions with the following free Ca^2+^ concentrations: 10^−10^ M Ca^2+^ (1 mM EDTA), 2 nM, 20 nM, 0.2 μM, 2 μM, 200 μM and 2 mM Ca^2+^. Due technical limitations, the measurements at free Ca^2+^ concentrations of 2 nM and 20 nM and 0.2 μM, 2 μM, 200 μM and 2 mM were carried out on different days. Control measurements in 1 mM EDTA solutions were carried out on both days (Fig 2A and B). The CMI1 CD spectra at free Ca^2+^ concentrations ranging between 0.2 μM to 2 mM were similar but were all significantly different from the 1 mM EDTA Ca^2+^ -free solution (Fig 2A). The CD spectra of CMI1 in 20 nM free Ca^2+^ concentrations were also significantly different from the Ca^2+^ free 1 mM EDTA solution and also different spectra were observed at 2 nM free Ca^2+^ (Fig 2B). The percentage of α-helix that were calculated based on the CD spectra were around 40% for free Ca^2+^ concentrations ranging between 0.2 μM to 2 mM and below 30% for CMI1 in the Ca^2+^-free 1 mM EDTA solution (Fig 2C). While the percentage of α-helix (Fig 2D) were lower compared to the measurements presented in panel (2C), the differences in α-helix content between the 20 nM free Ca^2+^ and 1 mM EDTA were around 10%, similar to the differences between the 0.2 μM-2 mM Ca^2+^ and the Ca^2+^-free 1 mM EDTA solutions (Compare Fig 2C and D). The CD spec analysis suggested that CMI1 can bind Ca^2+^ at free Ca^2+^ concentrations ranging between 10^−9^-10^−8^ M, which in turn induce secondary structure changes that result in an increase in a-helicity. Using the R-GECO Ca^2+^ sensor, it has recently been reported that the resting cytoplasmic Ca^2+^ concentrations [Ca^2+^]_cys_ along the root range between 50-90 nM (35). Moreover, as indicated in figure 1, different cells types in the root appear to display specific and distinct resting Ca^2+^ concentrations. Thus, it appears well conceivable that CMI1 serves as a highly sensitive sensor already responding to minor fluctuations in Ca^2+^ concentrations and that CMI1 exerts its function in Ca^2+^ associated status.

**Fig 2.**
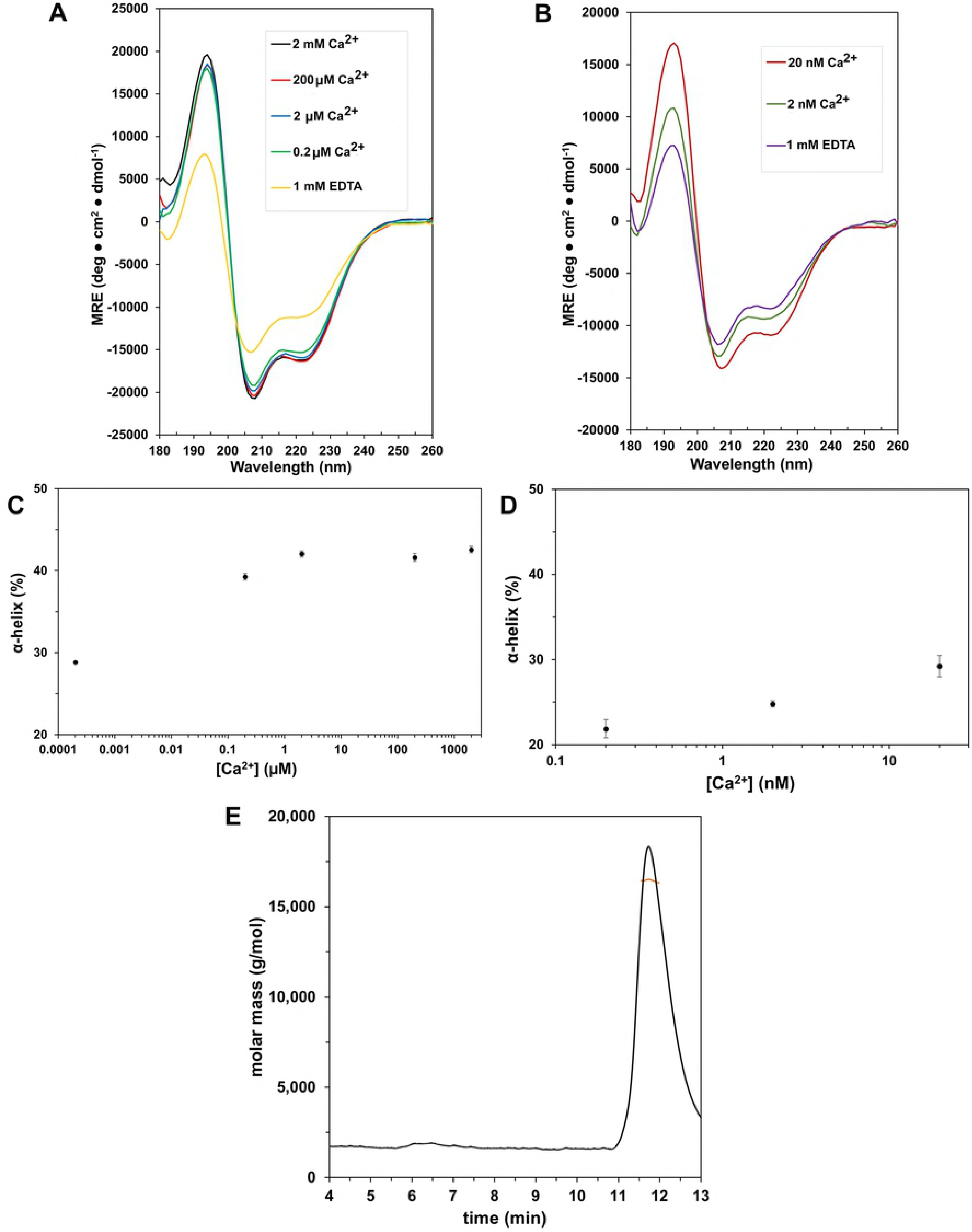
CMI1 changes secondary structure in free Ca^2+^ concentration ranging between 10^−9^ - 10^−8^ M and is a monomer in solution. (A and B) CD-spectra of 60 μM CMI1 at indicated free Ca^2+^ concentrations. Each curve is labeled as per legends. Measurements presented in panel A and B were carried out on different days. (C and D) Percent of α-helix of CMI1 at different free Ca^2+^ concentrations calculated from the CD spectra in A and B, respectively. (E) A SEC-MALS elution profile of 4 μg CMI1 in 2 mM Ca^2+^ solution. CMI1 eluted as a single peak with a molecular mass (red line) corresponding to a monomeric form.

Many Ca^2+^ binding proteins require at least two EF hands for their function or function as dimers if they contain an non-even number of EF hands. We therefore hypothesized that CMI1, bearing only a single Ca^2+^-binding EF-hand, might oligomerize in solution. Therefore, the quaternary structure of CMI1 was examined by Size Exclusion Chromatography Multi Angle Light Scattering (SEC-MALS). To eliminate potential effects of the poly-His-tag on solution structure, we performed the analysis on recombinant bacterially expressed and purified recombinant CMI1 from which the poly-His-tag was cleaved. At both concentrations of 2 and 4 mg/ml, CMI1 eluted as a monodisperse species at around 11.5 min with a measured molecular mass of 16 kDa, corresponding to the monomeric form of the protein (Fig 2E and S2 Fig). Hence, we conclude that CMI1 is strictly monomeric in vitro at least to concentrations of 25 μM in high Ca^2+^ conditions. This finding however does not exclude that CMI1 upon interaction with additional proteins may form oligomeric assemblies.

Possible homo- and hetero-dimerization of CMI1 was also examined by yeast two-hybrid assays. The analysis was carried out with Clontech^®^ LexA yeast two hybrid yeast strain *EGY48*, since CMI1 activates gene expression in Gal4-based yeast-2-hybrid strains when expressed fused to Gal4 DNA binding domain (Gal4-DB). Following 24 hours incubation, very faint blue color appeared in X-Gal assays of CMI1-LexA-BD/CMI1-Lex-AD (activation domain) compared to strong blue color in the CMI1-ICR1 and no color in the vector-control assays. Following 48 hours incubation, the X-Gal assays of the CMI1-BD/CMI-AD assays had light blue color compared strong blue of the CMI1-ICR1 and no color in the negative vector control assays (S3 Fig). Together, the yeast two-hybrid assays suggest that CMI1 could form dimers in yeast in the absence of ICR1 but also that the high affinity to ICR1 would interfere with this homo-dimerization. Therefore, the differences in the strength of the interaction in yeast and the SEC-MALS results strongly suggest that CMI1 very likely interacts with ICR1 as a monomer.

### Interaction of CMI1 with ICR1 involves a conserved hydrophobic pocket in CMI1 and a Calmodulin (CaM) Binding-like Domain (CBLD) in ICR1

Having established that CMI1 could function as a Ca^2+^ sensor we sought to obtain more insights into the molecular details of its structure and function. The 3D structure of CMI1 was predicted using homology modeling based on the structure of KIC, which belongs to the same subfamily single EF-hand Ca^2+^ binding proteins (32, 36). The predicted structure of CMI1 suggests the formation of two helix-loop-helix domains, one which binds Ca^2+^ and one which does not. This structural feature likely enables CMI1 to function as a monomer with regard to Ca^2+^ binding (Fig 3A and B). The CMI1 structure with Ca^2+^ bound is predicted to form a hydrophobic pocket (Fig 3A, residues highlighted in yellow). Modeling of CMI1 in complex with the Calmodulin (CaM) Binding Domain (CBD) of the KIC interactor Kinesin-like Calmodulin Binding Protein (KCBP) (32, 36) revealed that three Leu residue in the putative hydrophobic pocket of CMI1 namely L59, L92 and L100 can potentially serve as interacting side chains with a Trp residue in a domain that would be structurally related to a CBD which we therefore designated as Calmodulin (CaM) Binding-like Domain (CBLD) (Fig 3B).

**Fig 3.**
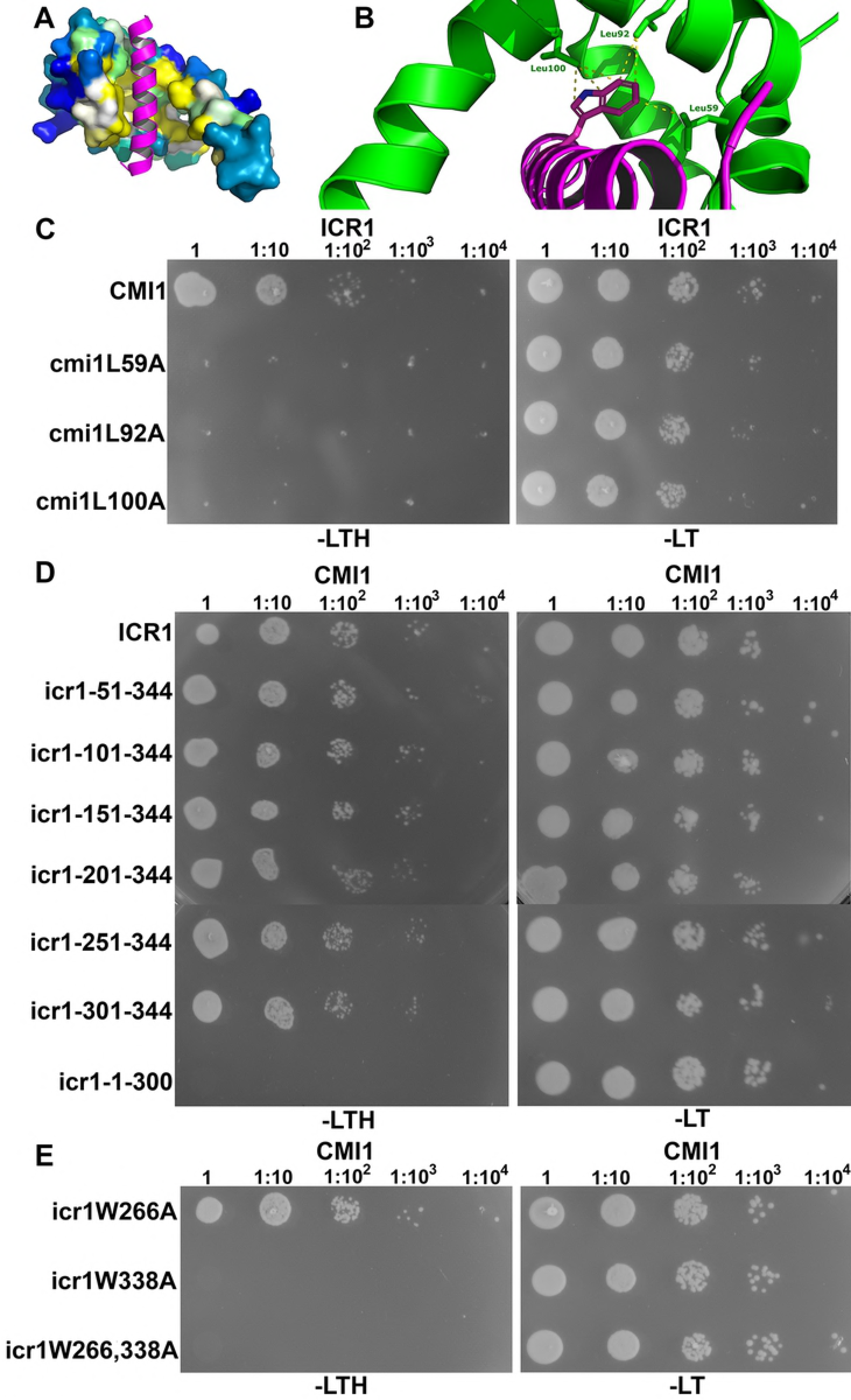
Interaction between CMI1 and ICR1 requires a hydrophobic pocket in CMI1 and a C-terminal W338 residue in ICR1. (A and B) A homology model of CMI1 with the Calmodulin Binding Domain (CBD) of KCBP shown in magenta. (A) A surface representation of CMI1 with residues of the hydrophobic pocket highlighted in yellow. (B) A closeup displaying CMI1 Leu residues L59, 92 and 100 (green) interacting with a Trp residue in KCBP CBD (magenta). (C-E) Yeast 2-hybrid assays. (C) ICR1 did not interact with CMI1 hydrophobic pocket L59, L92 and L100 mutants. (D) ICR1 44 C-terminal residues are required and sufficient for interaction with CMI1 but interactions are detected also at 1:10^4^ dilution with icr1-151-344 C-terminal or longer fragments. (E) ICR1 Trp residue W338 but not W266 is required for the interaction between CMI1 and ICR1. -LT: Leu, Trp deficient medium; -LTH: Leu, Trp, His deficient medium. (C-E) Numbers above panels denote dilutions of the yeast cells.

To test the hypothesis that L59, L92 and L100 form a hydrophobic pocket, we exchanged L to A in each of the respective Leu residues and tested the interaction of this modified CMI1 versions with ICR1 in yeast. As expected, neither cmi1 mutants L59A, L92A nor L100A interacted with ICR1 in yeast two hybrid assays (Fig 3C), strongly suggesting that the three Leu residues are part of a hydrophobic pocket required for protein-protein interaction.

There are two Trp residues in the C-terminal end of ICR1 at positions 266 and 338 that could be part of a potential CBLD. To map a potential CMI1-interaction domain in ICR1, we generated a series of N and C-terminal deletion mutants of ICR1 and examined their interaction with CMI1 in yeast two hybrid assays. These analyses revealed that the 44 C-terminal residues of ICR1 (icr1-301-344) are necessary and sufficient for interaction with CMI1 (Fig 3D). Slightly stronger yeast growth that resembled the full-length ICR1 was observed between an ICR1 C-terminal fragment encompassing residues 151 to 344 (icr1-151-344) and CMI1 (Fig 3D). Hence, the C-terminal 44 residue domain of ICR1 can function as a CBD but possibly other residues also contribute to the interaction.

Next, we examined the interaction between CMI1 and ICR1 harboring the single amino acid substitutions W266A and W338A. In yeast two hybrid assays, ICR1W266A still interacted with CMI1 while ICR1W338A did not (Fig 3E). Similar results were obtained when W266 and 338 were mutation to Gln (Q) (S4 Fig). Together, these results suggest that W338 is the primary Trp residue in ICR1 CBLD that is most crucial for interaction with residues in the hydrophobic pocket of CMI1.

Next, we examined the localization of CMI1, ICR1 and the potential influence of interaction with ICR1 on CMI1 localization in plants. When expressed in plants, ICR1-mCherry localized to microtubules (MTs) as indicated by its colocalization with the MTs marker TUA6-GFP (S5A-C Fig). When expressed by itself in *Arabidopsis* under control of its own promoter CMI1 was observed at the plasma membrane, throughout the cytoplasm and in nuclei (S6 Fig). Imaging of leaf epidermis pavement cells showed the mRFP-CMI1 is indeed localized to the plasma membrane as well as to nuclei and cytoplasm (S6A Fig). Furthermore, protein immunoblot with anti-CMI1 antibodies indicated that CMI1 is localized in soluble and insoluble fractions in different tissues (S6B and C Fig). However, when ICR1 and CMI1 were transiently coexpressed in *N. benthamiana* leaf epidermis cells, ICR1-mCherry and GFP-CMI1 were localized to MTs (Fig 4A-C). The colocalization of both mCherry-ICR1 and GFP-CMI1 was sensitive to the anti-MT drug oryzalin, confirming that they were both localized to MTs (S5D-F Fig). In contrast to GFP-CMI1, neither GFP-CMI1D85N (mutated in the Ca^2+^ binding EF hand) nor GFP-CMI1L59A (mutated in the hydrophobic pocket) were recruited to MTs (Fig 4D-F and G-I). Likewise, when GFP-CMI1 was coexpressed with ICR1W338A-mCherry it was not recruited to MTs while ICR1W338A-mCherry was observed on MTs (Fig 4J-L). Taken together, the coexpression assays in plants reinforced the combined conclusions derived from the results of the structural modeling and interaction assays, demonstrating that also in plant cells the interaction between ICR1 and CMI1 is Ca^2+^-dependent, involves a hydrophobic pocket in CMI1 and a C-terminal CBLD involving W338 in ICR1. These results also provide the opportunity that CMI1 modulates the function of ICR1 and/or fulfills alternative functions in its ICR-bound and ICR-non-bound form.

**Fig 4.**
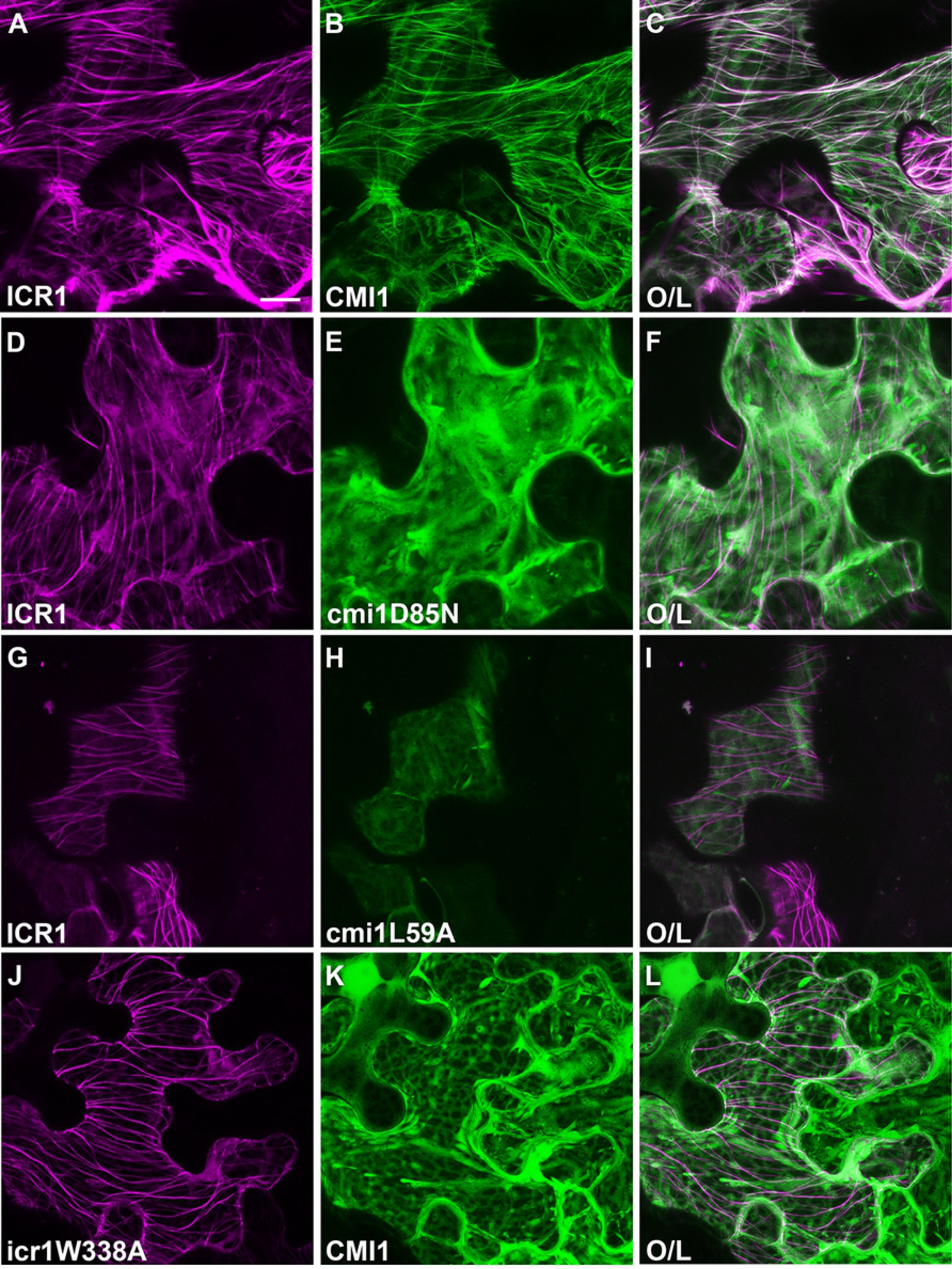
Recruitment of CMI1 by ICR1 to MTs depends on Ca^2+^ binding and intact hydrophobic pocket of CMI1 and ICR1 W338. (A-I) CMI1 but not Ca^2+^ non-binding cmi1D85N and hydrophobic pocket cmi1L59A mutants is recruited to MTs by ICR1. (J-L) icr1W338A is associated with MTs but does not recruit CMI1. Each panel is as per legends. O/L-overlay of mCherry and GFP signals. Bar, 20 μm for all panels.

### Expression of CMI1 is regulated by auxin through TIR1/AFB auxin receptors

To gain first indications for the function of CMI1 in plants, we examined the expression pattern and regulation of CMI1 expression and their correlation with cytoplasmic Ca^2+^ levels. High CMI1-GUS and mRFP-CMI1 levels were detected in the root meristem and lateral root primordia of *pCMI1::CMI1-GUS* and pCMI1>>mRFP-CMI1 plants (Fig. 5A and B and S7A Fig), resembling the expression pattern of *DR5* promoter driven auxin reporters (37). Regions of increased expression of CMI1-GUS in the root elongation and maturation differentiation zones of *pCMI1::CMI1-GUS* plants were observed following treatments with 10 μM IAA (Fig. 5C and D), resembling the pattern of TIR/AFB auxin induced genes (26). A qPCR analysis confirmed induction of CMI1 mRNA following auxin treatments (S7B Fig). In agreement, microarray experiments revealed that induction of *CMI1* expression by auxin was reduced in the *axr1* (auxin resistant 1) auxin signaling mutant (38), indicating that auxin induces *CMI1* expression by a TIR1/AFB dependent mechanism (Fig 5E). Furthermore, our analysis of additional publicly available microarray data revealed that *CMI1* was induced by exogenous auxin treatments and suppressed in the *axr2-1/iaa7* auxin insensitive mutant (39). Taken together, these results indicate that the expression of *CMI1* is enhanced in cells and tissue with increased auxin concentration and also regulated by auxin via the TIR1/AFB auxin receptor system.

**Fig 5.**
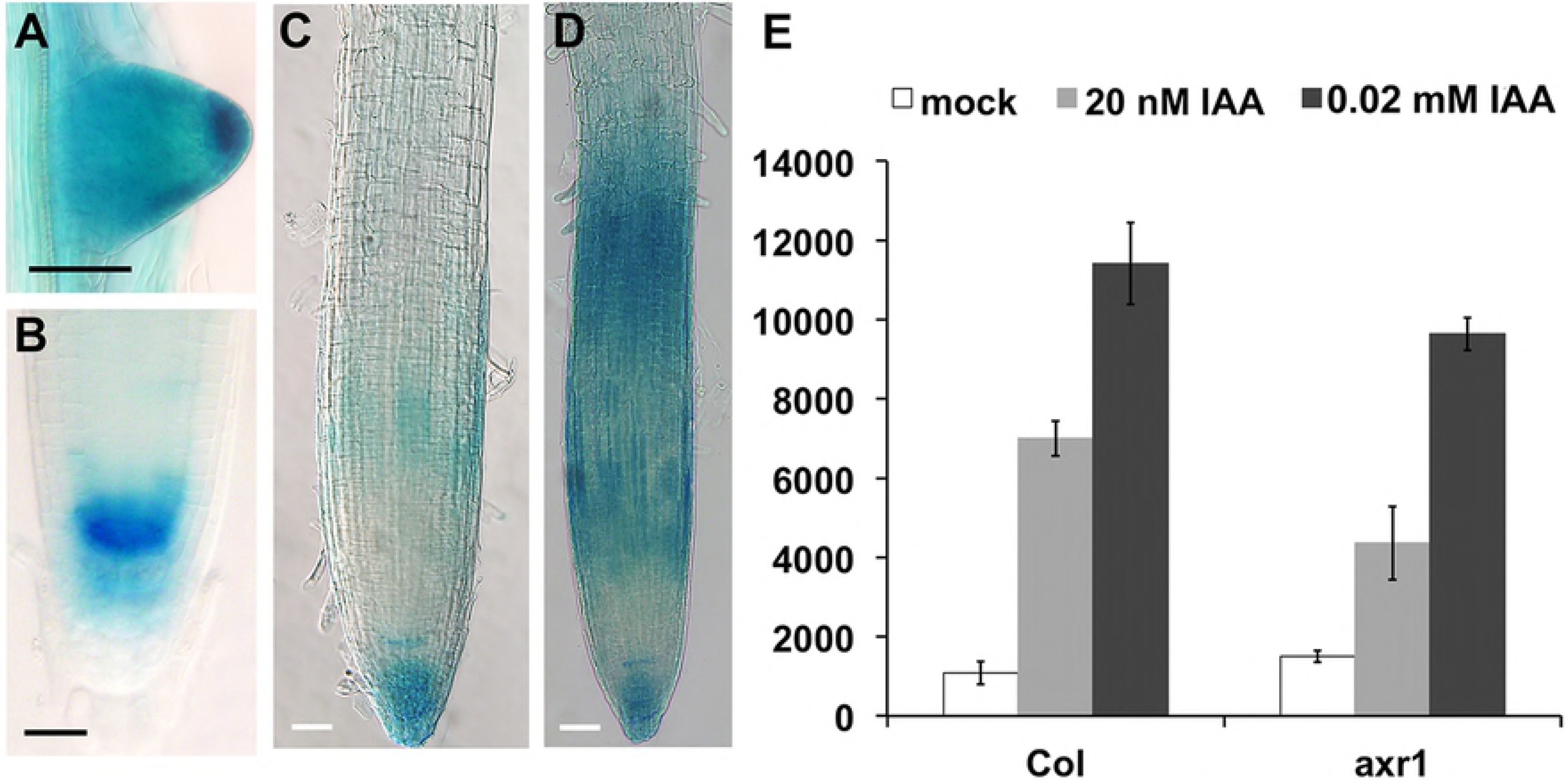
Expression of CMI1 is induce by auxin via TRI/AFB receptors. Expression of CMI1-GUS in lateral root initial (A) and primary root meristem and (B). (C and D) Expression level and pattern of *pCMI1* driven CMI1-GUS in *cmi1* mutant background without (C) and 2 hours following treatment with 10 μM IAA (D). (E) Microarray expression data showing the induction of *CMI1* by auxin is reduced in *axr1* auxin response mutant background. Scale bars, 20 μm.

### CMI1 mediates auxin responses and fine tunes root growth

Next, we further examined the function of CMI1 and its interconnection with auxin signaling in plants by analyzing the phenotype of a *CMI1* loss of function mutant. The *cmi1* mutant (Cold Spring Harbor Laboratory (CSHL) line GT_24505) carries a transposon insertion at nucleotide 37 in the *CMI1* coding region (Fig 6A, B). Compared to a wild type control, the *cmi1* plants have shorter primary roots (Fig 6C, D) as a result of a smaller root meristem size, defined as the length of the region between the QC and the initiation of the elongation zone (Fig 6E, F). Importantly, the shorter primary root phenotype was complemented by *pCMI1::CMI1-GUS* (S8A Fig), confirming that the mutant phenotype resulted from the loss of *CMI1* function and that the observed expression pattern of *pCMI1::CMI1-GUS* reflects the expression pattern of the endogenous *CMI1* gene.

**Fig 6.**
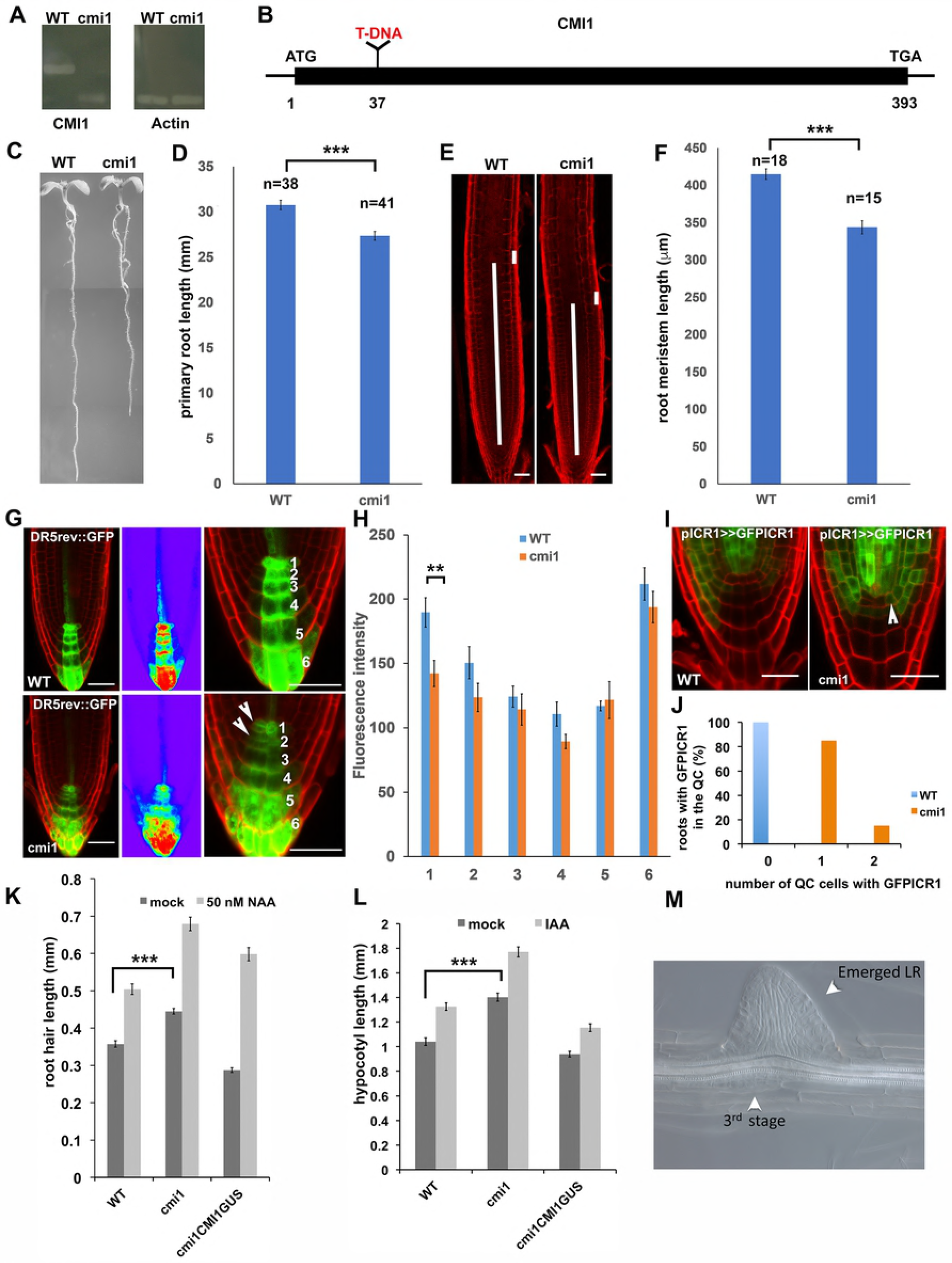
*cmi1* mutant plants have higher ICR1 levels in the QC and auxin-related phenotypes. (A) The *CMI1* RNA cannot be amplified in *cmi1* indicating that the mutant is a null. (B) A diagram of the *CMI1* gene highlighting the T-DNA insertion at position 37. (C) 7-days old *cmi1* seedlings have shorter primary roots. (D) Quantification of the root length in *WT*(*Ler*) and *cmi1* plants. Error bars are SE, p≤0.001 (T-test). (E) Root cell division zones of WT (*Ler*) and *cmi1* 7-days old seedlings. The long bars highlight the measured root zone length. The short bars show the cell length used to determine the end of the cell division zone. (F) Quantification of the root cell division zone length calculated with root samples as shown in panel E. Error bars are SE, p=6.42×10^−7^ (p≤0.001), T-test. (G and H) DR5_rev_::GFP auxin response maximum is reduced in *cmi1* QC. (G) Cell walls were stained with PI. The middle panels show heat diagram of the roots shown in the left panels. Right panels show higher magnifications used for quantifications. The numbers correspond to cell layers. Arrowheads highlight the signal reduction in *cmi1* compared to WT. (H) Quantification of DR5_rev_::GFP fluorescence intensity in cell layers 1-6 as defined in panel G. Layer 1 is the QC. Error bars are SE, p=0.006 (p≤0.01), T-test. (I) GFP-ICR1 expression is up-regulated in the QC (arrowhead) in *cmi1* roots. (J) Percentage of WT and *cmi1* roots with GFP-ICR1 expression in 1 or 2 QC cells. (K) Root hair length in *Ler* (WT), *cmi1* and *cmi1* complemented with CMI1-GUS (cmi1CMI1GUS) in control (mock) or following treatments with 50 nM NAA. The root hairs in *cmi1* mutants are significantly longer than in the wild type and CMI1-GUS complemented roots. Bars are SE, p≤0.001 (T-test). (L) Hypocotyl length is increased in *cmi1* mutants and in response to 5 μM IAA treatments. hypocotyls of *cmi1* mutants are significantly longer than the wild type and CMI1-GUS complemented seedlings. Bars are SE, (p≤0.001, T-test). (M) A stage 3 LRI developing opposite to an emerging LRI in a *cmi1* root. Scale bars, 50 μm in E and 20 μm in G, I.

Next, we determined whether the loss of *CMI1* function affects auxin distribution using the *DR5*_*rev*_::*GFP* auxin response marker (40, 41). In *cmi1* mutant roots, the auxin response maximum in the QC was reduced, compared to a wild type control (Fig 6G). Quantification of the GFP fluorescence levels revealed a significant reduction (p≤0.006, T-test) in fluorescence level in the QC cells (Fig 6H). Ectopic expression of GFP-ICR1 was detected in the QC cells of the *cmi1* mutant, but not in wild type, roots (Fig 6I, J), in line with the reduced auxin response in the QC (30, 31). Hence, CMI1 affects ICR1 levels, indirectly by regulating the auxin response.

The regulation of *CMI1* expression by auxin prompted us to examine the possible involvement of CMI1 in further well characterized auxin responses. The initiation and elongation of root hairs are regulated by TIR1/AFB-Aux/IAA-dependent auxin signaling (42–44). We found that root hairs were longer in the *cmi1* mutant, compared to wild type and *cmi1/pCMI1::CMI1GUS* plants, and elongated in response to exogenous auxin treatments (Fig 6K and S8C Fig). TIR1/AFB-AUX/IAA dependent auxin signaling also affects hypocotyl length (45, 46). Hypocotyls of the *cmi1* mutant were significantly longer (p≤0.001, T-test), compared to wild type and *cmi1/pCMI1::CMI1GUS* plants. As expected, external IAA treatments induced hypocotyl elongation in wild type, *cmi1* mutant and *cmi1/pCMI1::CMI1 GUS* plants (Fig 6L and S8B Fig). Lateral root formation is regulated by both auxin response and distribution (40, 47–50). *cmi1* plants exhibited abnormal lateral root patterning (Fig 6M and S9 Fig) with an average of 7 LRs/cm in *cmi1* compared to 4 LR/cm in control *Ler* seedlings. Together, the changes in DR5::GFP_rev_ and GFP-ICR1 expression pattern and the macroscopic phenotype of *cmi1* mutant plants suggest that CMI1 regulates both the spatial distribution and the level of auxin responses.

Corresponding to the increased auxin response of *cmi1* mutants, the *DR5::GUS* staining was stronger in *cmi1* primary root and lateral root initials compared to wild type control (Fig 7A-J). To further examine the function of CMI1, we ectopically expressed mRFP-CMI1 under regulation of the ICR1 promoter (*pICR1>>mRFPCMI1*), using a transcription transactivation system (51). The roots of *pICR1>>mRFPCMI1* plants were short, had reduced columella layers and reduced auxin response maxima (Fig 8A-F). Hence, ectopic expression of CMI1 was associated with repression of auxin responses and root growth.

**Fig 7.**
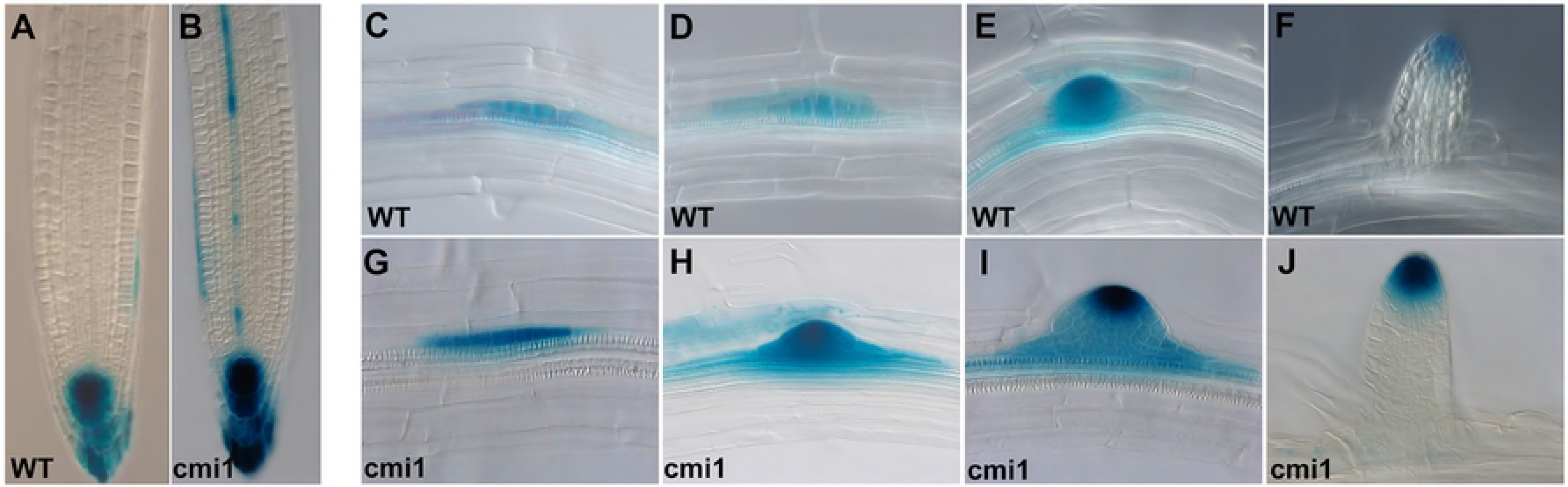
CMI1 loss of function results in enhanced auxin induced DR5::GUS expression. (A) Expression level of *DR5::GUS* auxin response marker in roots of *L. erecta* (WT) and *cmi1* (B). (C-J) Expression levels of *DR5::GUS* in LRI of Ler (WT) (C-F) and *cmi1* (G-J).

**Fig 8.**
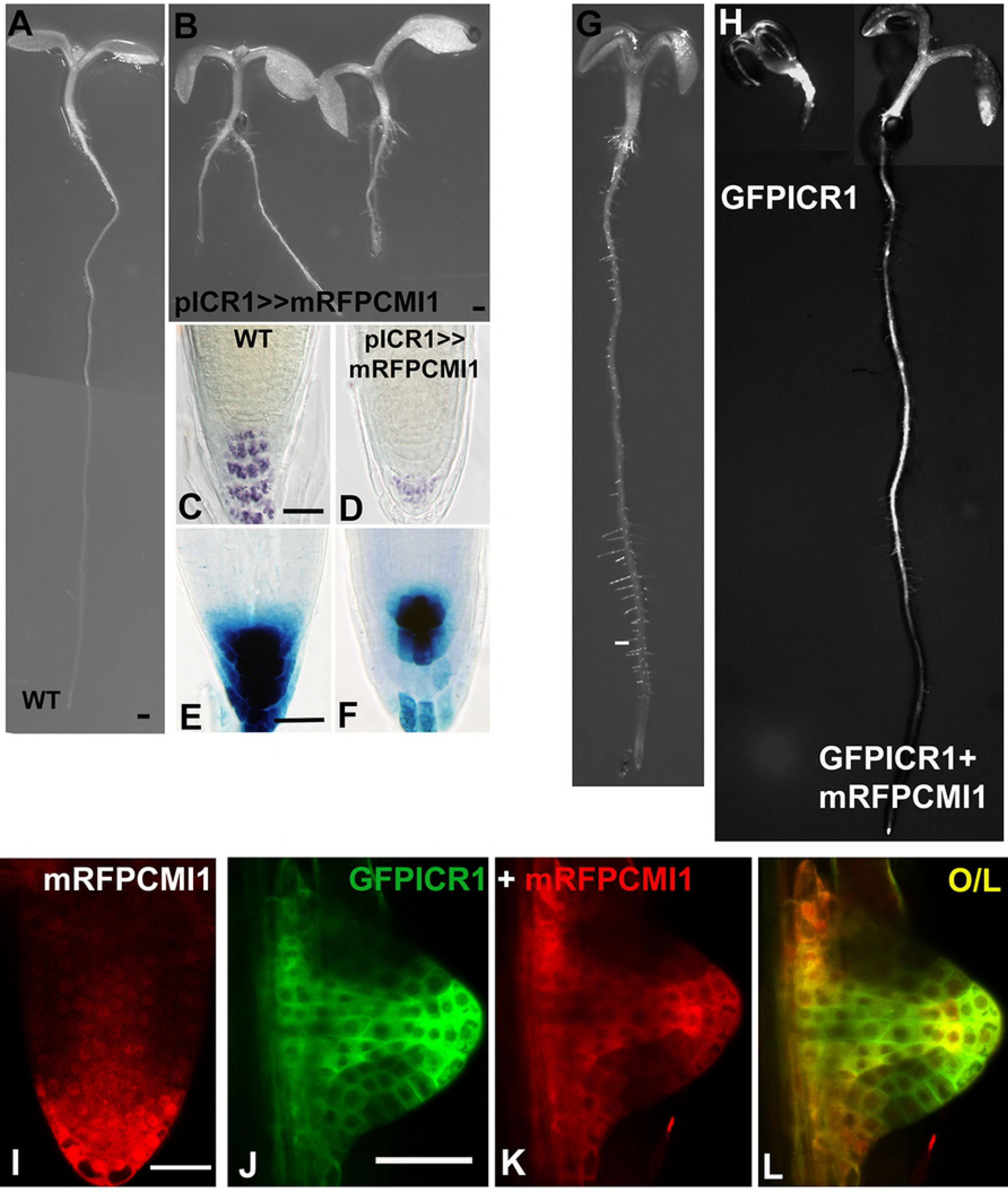
Ectopic expression of CMI1 suppresses root development and auxin response. (A) Control *Col-0* (WT) seedling. (B) Root growth is arrested in *pICR1>>mRFP-CMI1* seedlings. (C and D) Reduced iodine (IKI) columella staining in a *pICR1>>mRFP-CMI1* root. (E and F) Reduced auxin response in a *pICR1>>mRFP-CMI1 DR5::GUS* root. (G) A control pCMI2>>LhG4 plant. (H) Root development is inhibited in a pCMI2>>GFP-ICR1 plant (left) and restored by coexpression of GFP-ICR1 and mRFP-CMI1 in pCMI2>>GFP-ICR1/mRFP-CMI1 plants (right). (I) mRFP-CMI1 is expressed in the lateral root meristem QC and initial cells and 1133 accumulates in the cytoplasm and nuclei in pCMI1>>mRFP-CMI1 plants. (J-L) GFP-ICR1 and mRFP-CMI1 are colocalized in the cytoplasm in a pCMI2>>GFP-ICR1/mRFP-CMI1 lateral root initial. Note the absence of mRFP-CMI1 from nuclei. Scale bars 0.5 mm (A, B, G and H), 50 μm (C-F) and 50 μm (I-L).

Previously, we demonstrated that inducing elevated levels of ICR1 in the QC by its expression under regulation of the CMI2 promoter, utilizing the pOp/LhG4 transcription/transactivation system (pCMI2>>GFP-ICR1), resulted in inhibition of root growth (31) and (Fig 8G and H). Remarkably, co-expression of GFP-ICR1 and mRFP-CMI1 in pCMI2>>GFP-ICR1/mRFP-CMI1 resulted in suppression of root growth arrest (Fig 8H). In lateral root primordium mRFP-CMI1 was detected in nuclei, cytoplasm and plasma membrane (Fig 8I), similar to its distribution in the leaf epidermis pavement cells (S6A Fig) and in agreement with the protein immunoblot with anti CMI1 antibodies that indicated localization in both soluble and insoluble fractions (S6B Fig). Examination of the subcellular localization of both GFP-ICR1 and mRFP-CMI1 revealed that when co-expressed together with GFP-ICR1 in the same cells, mRFP-CMI1 was excluded from nuclei (Fig 8J-L). Together, these data indicated that similar to transient expression in *N. benthamiana*, co-expression of CMI1 and ICR1 in the same cells affected the subcellular distribution of CMI1 and its function.

To examine a potential impact of CMI1 function on the regulation of auxin transport and PIN polarity we carried out whole mount Immuno-staining of roots with anti PIN1 and PIN2 antibodies. Importantly, these immunostaining experiments revealed that PIN2 distribution in the cortex is altered in cmi1 (S10A Fig). In 55% of the cells PIN2 displayed apical localization and in another 25% it was non-polar. In comparison, in *Ler* (WT) in 90% of the cell PIN2 displayed basal localization in the cortex and in only 10% of the cells it was either apical or non-polar (S10B Fig). Previous studies have shown that the CMI1 closest homolog CMI2/PID Binding Protein 1(PBP1)) interacted with the AGCIII kinase PID, which regulates PIN polarity and function (24). Hence, we tested the interactions between CMI1 and PID as well as other know auxin signaling components in yeast two hybrid assays (S11 Fig). However, none of these auxin signaling associated proteins interacted with CMI1. Therefore, the clarification of the mechanism how CMI1 regulates auxin responses and PIN2 polarity awaits further experimental clarification in the future. Nevertheless, the phenotypic alterations upon perturbation of CMI1 function and their association with altered auxin responses and distribution unambiguously identify the Ca^2+^ sensor CMI1 as a critical component modulation the action of auxin as regulator of root growth and differentiation.

### CMI1 affects auxin induced Ca^2+^ signaling in a cell type/tissue specific manner

To further examine a potential role of CMI1 in interconnecting Ca^2+^ signaling and auxin function possibility, we compared the auxin-induced cytoplasmic Ca^2+^ signals in the Lateral Root Cap (LRC), epidermis and vasculature of wild type and *cmi1* roots expressing YC3.6. The quantification of Ca^2+^ dynamics was carried out by calculating the ratio change in the FRET signals (ΔR/R_0_) (Fig 9A-F). Quantitative analyses revealed genotype specific differences in the tissue specificity and intensity of auxin-induced cytoplasmic Ca^2+^ signals between *Ler* (WT) and *cmi1*. In the LRC, the auxin-induced cytoplasmic Ca^2+^ response displayed a higher amplitude in the wild type *Ler* than in *cmi1* mutants. Specifically, none of the *cmi1* roots, as opposed to 35% of the analyzed wild type roots, showed a ratio change ΔR/R_0_≥0.15. In contrast, 40% of the *cmi1* roots displayed lower Ca^2+^ elevations, characterized by ΔR/R_0_ ranging between 0-0.1, while only 5% of the wild type roots displayed such low Ca^2+^ signals in the LRC (Fig 9A and B). Additionally, the *cmi1* mutants exhibited a faster kinetics in restoring the basal level of Ca^2+^. In the epidermis, strong increases in Ca^2+^ levels (ΔR/R_0_≥0.15) predominated in both the wild type and *cmi1* backgrounds, with only slightly less samples with high Ca^2+^ levels detected in *cmi1* mutants (Fig. 9C and D). In the vasculature, the amplitude of the auxin-induced Ca^2+^ signals were comparable in wild type and *cmi1* mutant. The low threshold Ca^2+^ signals (ΔR/R_0_ ranging between 0-0.1) predominated in both wild type and *cmi1* backgrounds and only 10% more wild type roots displayed higher Ca^2+^ levels, with ΔR/R_0_ values ranging between 0.1-0.15 (Fig 9E and F). However, there was a striking difference in the shape of the signal and in the kinetics of the signal to reach the maximum amplitude. The wild type roots evoked a maximum Ca^2+^ response in 120 s, while in the cmi1 mutants the Ca^2+^ maxima were reached in 230s. Interestingly, in the vasculature the restoration of basal level of Ca^2+^ followed a similar kinetics. Taken together, these results reveal that loss of *CMI1* function alters auxin-induced Ca^2+^ signals, especially in the lateral root cap and vascular cells, and suggest that CMI1 regulate auxin-associated changes in cytoplasmic Ca^2+^ levels in a cell/tissue specific fashion. Moreover, our finding point to an elaborate cell specificity and diversity in complex tissues.

**Fig 9.**
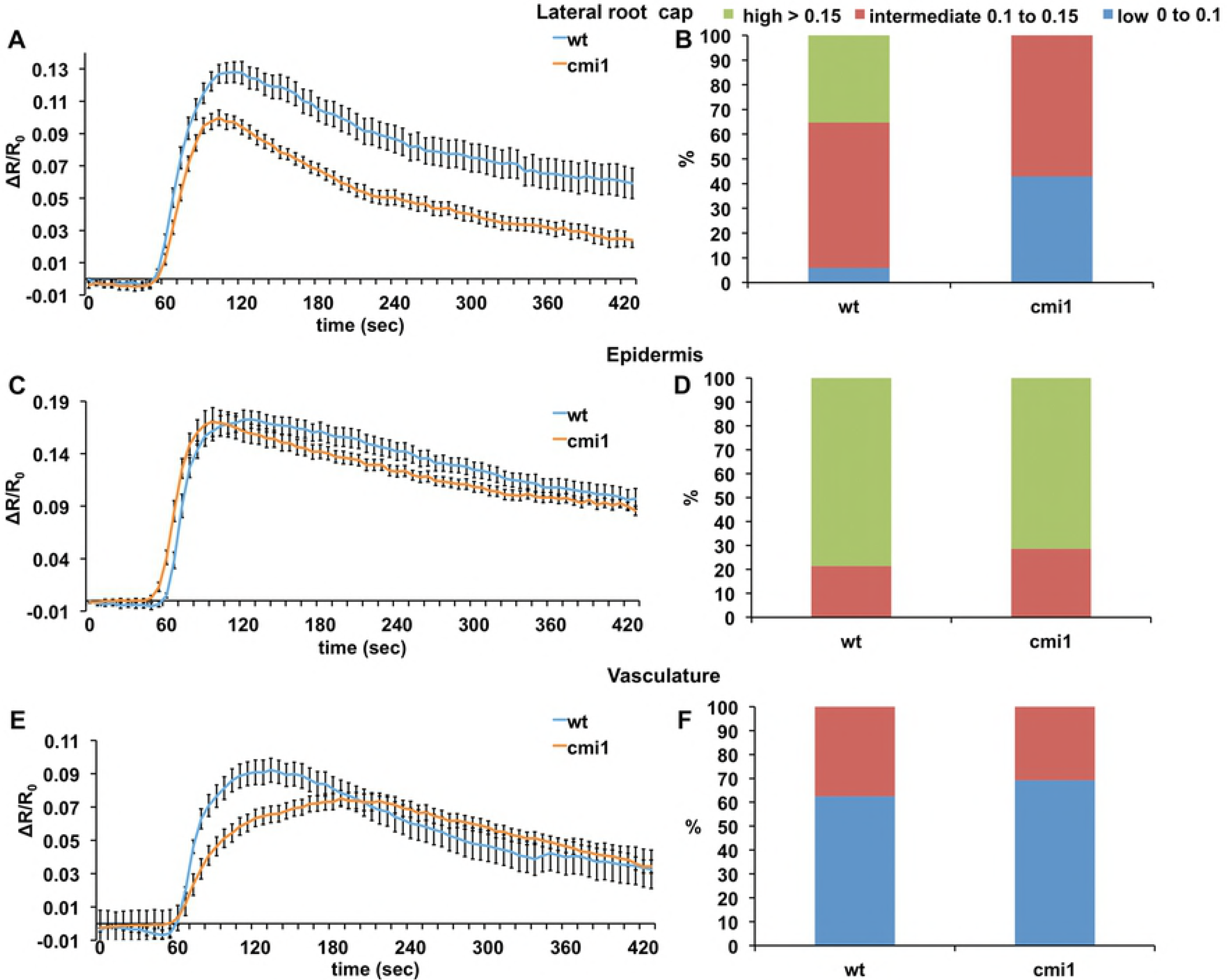
Auxin-induced Ca^2+^ response are reduced and display a different kinetic in *cmi1* in a tissue specific fashion. Auxin induced Ca^2+^ responses in root lateral root cap (A and B), epidermis (C and D) and vascular tissues (E and F). Note the reduced Ca^2+^ levels and different kinetics in Ca^2+^ decrease and increase in the lateral root cap and the vascular tissues, respectively.

## Discussion

The results presented in this work indicate that CMI1 serves as a Ca^2+^ sensor that links auxin and Ca^2+^ signaling. CMI1 expression is regulated by auxin, coincides with auxin induced cellular Ca^2+^ increases and the phenotype of *cmi1* mutant plants is associated with impaired auxin responses.

In yeast 2-hybrid assays, we did not detect interaction of CMI1 with components of the auxin transport and signaling machinery (Fig. S11). While the negative results of yeast 2-hybrid assays do not exclude interactions with these components they suggest that these interactions are unlikely. It is yet unclear whether CMI1 effects on auxin distribution and response involve its interaction with ICR1. CMI1 levels are highest in the QC where ICR1 is post-translationally degraded (30, 31), suggesting that its function in QC is different than in surrounding tissues. Whether and how CMI1 function in the QC is related to auxin accumulation in these cells is yet to be determined.

Despite having a single Ca^2+^-binding EF-hand, CMI1 most likely interacts with ICR1 and possibly with other target proteins as a monomer. The 3-D structure of CMI1 homolog KIC revealed existence of two EF-hands: a canonical Ca^2+^-that lacks essential residues for divalent ion binding. Upon Ca^2+^ binding both the Ca^2+^-binding and Ca^2+^ independent EF-hands form an open conformation, creating the hydrophobic pocket that can accommodate the KCBP CBD (36). Our structural modeling, structure-function assays and the SEC-MALS results indicate that like KIC, CMI1 has a canonical Ca^2+^ binding EF-hand and a Ca^2+^-independent EF hand which enable it to function as a monomer. Possibly, the oligomerization of ICR1, which contains a long coiled-coil domain (29) may induce accumulation of CMI1 molecules at discrete cellular domains.

The ability of CMI1 to responds to Ca^2+^ concentrations that are equal or lower than resting cellular Ca^2+^ levels suggest that it might not be regulated by fluctuations in cellular Ca^2+^ levels or that it is regulated by subtle change in cellular Ca^2+^ levels. The strong upregulation of CMI1 expression by auxin suggest that it transduces Ca^2+^-dependent responses auxin-dependently. The ICR1-modulated subcellular localization of CMI1 may suggest that CMI1 function is also regulated by its subcellular distribution and protein-protein interactions. The identification of the hydrophobic pocket in CMI1 and its interaction with ICR1 via a CBLD suggest that CMI1 may interact with other proteins which contain a CBLD. Given that the interaction of CMI1 with the C-terminal ICR1 CBLD is weaker than its interaction with longer fragments of ICR1, it is likely that the binding specificity between the proteins is determined by additional residues in ICR1. Thus, it is difficult to predict which CBLD containing proteins would interact with CMi1.

While KIC and CMI1 do not share common binding partners, it is interesting that they both interact with MTs binding proteins (this work and (32)). Unlike KCBP, which is a kinesin with enzymatic activity, ICR1 is a coiled-coil domain protein that does not contain additional known catalytic or structural domains and likely functions as a scaffold (29). KIC inhibits interaction of KCBP with MTs and its ATPase activity (32, 36). Data in this work indicates that ICR1 can recruit CMI1 to MTs, yet, it is unknown whether Unfortunately, in vitro assays to test the effect of CMI1 on ICR1 MT binding were unsuccessful because of the requirement to include Ca^2+^ in the reaction medium, which in vitro leads to MTs destabilization.

It has previously been shown that expression of a CMI1 homologue from wheat called TaCCD1 was induced be fungal elicitors (52). In rice, OsCCD1 was induced by ABA and osmotic stress and OsCCD1 overexpressing and mutant plants displayed increased and decreased salt tolerance, respectively (53). Publicly available transcriptomics data indicated that *CMI1* expression is repressed following 3 hours treatments with 140 mM NaCl and is then upregulated in the stele (http://dinnenylab.dpb.carnegiescience.edu/browser/query (54)) and that it is strongly upregulated by treatment with Rapid Alkalizing Factor 1 (RALF1) peptide (39). In agreement, the expression of pCMI1::GUS-CMI1 was down-regulated following 3 and 6 h treatments with 140 NaCl. Both GUS assays and qPCR analysis showed that treatments with RALF1 induced rapid and transient 30-50 fold increase in *CMI1* RNA levels, which reached a peak after 15 minutes and return to basal levels after 4 hours (S12 Fig). RALF1 is a ligand of the receptor-like kinase FERONIA (FER), which has been implicated in cell wall sensing and immune responses (55). The expression data suggest that CMI1 could be part of stress induced gene expression and cell wall sensing mechanisms. Our results suggest that CMI1 may have multiple functions. Under steady state condition its expression is primarily regulated by auxin and it is involved in regulation of auxin responses or distribution. Biotic and possibly other stress conditions that affect the cell wall induce rapid and transient upregulation of CMI1 that in turn may transduce rapid Ca^2+^ dependent response even at low cellular Ca^2+^ levels. An exciting hypothesis is that CMI1 may function as an integrator of auxin and various stress responses. Under non-stress conditions CMI1 functions in fine tuning of auxin responses. Under biotic or other stimuli that elicit increase in RALF levels CMI1 is rapidly upregulated and in turn it may suppress auxin levels/responses. Salt stress induces transient changes in root elongation zone cells’ which have been associated with FER signaling dependent Ca^2+^ spikes along the root (56). Hence, transient down-regulation of CMI1 under salt stress conditions may be part of the response that enables recovery from the stress.

A tight link between TIR/AFB-dependent auxin signaling and short and long distance increases in [Ca^2+^]_cyt_ has recently been demonstrated (7). Our work, identified CMI1 which expression is developmentally tightly regulated by auxin and in turn it regulates, auxin responses and distribution and auxin induced changes in Ca^2+^ levels. The rapid expression regulation of CMI1 by different environmental stimuli together with the phenotype of CMI1 loss of function and overexpression implicate its functioning as an Ca^2+^ sensor integrator of auxin and stress stimuli.

## Materials and methods

### Molecular cloning

The Plasmids used in this study are listed in supplemental information S1 Table. *pICR1>>GFP-ICR1* and *pCMI2>>GFP-ICR1* plants were previously described (30, 31). *pCMI1::CMI1-GUS* (*pSY1804*) was constructed by amplifying a 2,526 bp fragment containing the 2040 bp promoter, 78 bp 5’-UTR and the 408 bp CMI1 ORF, in which the TGA stop codon was changed to TAA (Leu). The resulting fragment was digested with *EcoRI* and *SalI* and cloned into *pENTRY1a*. The resulting plasmid *pSY1802* was recombined with *pMDC162* using LR clonase to obtain *pSY1804. pCambia2300-RFP-CMI1* (pSY1351) was generated by cloning mRFP upstream to the CMI1 ORF into *pCambia2300*. Transactivation CMI1 promoter plasmid (*pSY1806*) was constructed as follows: a 2040 bp fragment of the *CMI1* promoter was amplified, digested with *Sal1* and subcloned into *pLhG4Bj36* upstream of the chimeric transcription factor LhG4 to create plasmid *pSY1805. pSY1805* was then digested with *NotI* and the resulting fragment containing *pCMI1::LhG4-terminator* was subcloned into *pART27* plant binary plasmid to obtain *pSY1806*. To obtain the *mRFP-CMI1 Op* reporter plasmid, the *mRFP-CMI1* fragment from *pSY1351* was digested with *HindIII* and *XhoI* and subcloned into *pOp* to obtain *pSY1807*. Subsequently, *pSY1807* was digested with *NotI* and the resulting fragment containing *10XOp::mRFP-CMI1* was subcloned into the plant binary vector *pMLBART* to obtain *pSY1808*. pGAD-CMI1 was created as follows: the coding sequence of CMI1 was amplified from cDNA and subcloned into pGET (Thermo Fisher Scientific). It was then digested with *BamHI* and *SalI* and the resulting fragment was ligated to pGAD vector to obtain *pSY1565*.

The generation of plasmids for yeast 2-hybrid and plant colocalization assays of site directed and deletion mutants of CMI1 and ICR1 and plasmids for expression of CMI1 in *E.coli* were carried out as follows. For site directed mutagenesis (SDM primers were designed using the QuikChange Primer Design tool found at Agilent web site (https://www.genomics.agilent.com/primerDesignProgram.jsp). SDM was perform with *Pfu-Ultra* DNA polymerase (S2 Table) followed by digestion with *DpnI* (S2 Table) to eliminate unwanted templates. In two cases, that the SDM did not provide the desired mutants (cmi1^L92A^), an alternative approach of a three-step overlap extension PCR reaction using Phusion DNA polymerase (S2 Table) was performed. From this point, the cloning steps were the same as described below.

Genes of interest were cloned with flanking ends of attB1/2 recombination sites using a two-step reaction of Phusion high-fidelity DNA polymerase (S2 Table). in cases that several DNA fragments were observed in the PCR reaction products, the relevant band was extracted using QIAEX II gel extraction kit (QIAGEN) or Wizard SV Gel and PCR Clean-Up System (Promega) (S2 Table). The attB1/2 flanking genes were transferred into *pDONR221* using the BP clonase reaction (S2 Table). All clones were verified by sequencing.

For yeast 2-hybrid, constructs were transferred by recombination from *pEntry221* and then by recombination to *pDEST22* (prey) or *pDEST32* (bait) using the LR clonase (S2 Table). Bait and prey plasmids were transformed into *PJ69-4a* yeast strain. Presence of respective plasmids was verified by yeast colony PCR (S2 Table).

For colocalization assays in plants, *CMI1*, *cmi1*^*L59A*^, *cmi1*^*D85N*^ were transferred by recombination from *pEntry221* to *pGWB6-35S::eGFp* using the LR clonase (S2 Table). In addition, 3-way GATEWAY recombination reactions (S2 Table) were carried out with *pEntryP4-P1R-35S* promoter, *pEntry221-ICR1 or pEntry221-icr1*^*W338A*^ (both without stop codon) and *pEntryP2R-P3-mCherry* into pB7m34GW. Plasmids were verified by colony PCR (S2 Table) and sequencing. For expression in plants, plasmids were transformed into *Agrobacterium tumefaciens* stain *GV3101 pMP90*.

Cloning for protein expression in *E. coli*. A PCR product of CMI1 with flanking BamH1 and Not1 sites was subcloned into pJET1.2 using the CloneJET PCR cloning kit (S2 Table). The resulting plasmid was digested with BamH1 and Not1 and the CMI1 fragment was subcloned into *pET21d.H8.Nia.yBRFc.T.GSTrc* digested also with BamH1 and Not1 to isolate the *pET21d-His*_*8*_-*TEV* fragment. The resulting plasmid *pSY2408 (pET21d_His8-TEV-CMI1*) was designed to express His_8_-TEV-CMI1 fusion protein that enables purification of CMI1 on a metal chelate Ni-column and cleavage of the His_8_-tag by TEV protease.

### Plant material and growth conditions

The *Arabidopsis* transgenic lines used in this study are listed in supplemental information S3 Table. Long-day grown (16 hours light/8 hours dark, 22° C) *Arabidopsis Columbia-0 (Col-0*) and *Landsberg erecta* (*Ler*) ecotypes were used for stable expression, mutant phenotypic analysis, protein localization and Ca^2+^ measurement. *Arabidopsis, mRFP-CMI1* and *pCMI1>>mRFP-CMI1* and *pICR1>>mRFP-CMI1* plants were generated by crossing, *pCMI1-LhG4* to *pop-mRFP-CMI1* and *pICR1-LhG4* to *pop-mRFP-CMI1*. The *cmi1* mutant (Cold Spring Harbor laboratory CSHL_GT24505) is in the *Ler* background. To analyze *DR5::GFp_rev_* and *pICR1>>GFP-ICR1* expression in the *cmi1* mutant background, *DR5::GFp_rev_* and *pICR1>>GFP-ICR1* were crossed into wild type *Ler* and *cmi1* backgrounds. M3 generation wild type or *cmi1* homozygote mutant plants that harbored the *erecta* phenotype and expressed either *DR5::GFp_rev_* or *pICR1>>GFP-ICR1* were selected for the further analysis. Quantification of fluorescent signals was performed using Image J. For *DR5::GFPrev* quantification we used 1216 images of independent root tips when 2-4 QC cells are in the center. Cell layers 1-6 were defined from QC to the last columella layer and GFP signal intensity was measured in the same area (below the QC cells) in each layer using Image J. The average of the GFP intensity is presented in the graph and the bars are the SE (Fig 6). To quantify the ectopic expression of GFP-ICR1 in the QC cells of *cmi1* mutant, 18-20 root of each WT (Ler) or *cmi1* plants were imaged when QC cells (2-4 cells) are visible in the center. The number of QC cells, in which a GFP-ICR1 signal was detected, was used to calculate the percentage of the roots with or without ectopic expression. Complementation of *cmi1* was performed by crosses with *pCMI1::CMI1-GUS* plants. The analysis was performed using non-segregating lines from the fourth and fifth generations. For Ca^2+^ imaging the *pUBQ10::YC3.6* Yellow Cameleon (25) was transformed into *Ler* wild type and *cmi1* plants. Several independent transgenic lines were used for the Ca^2+^ imaging.

### Protein expression and antibody generation

Expression in *E. coli* Rosetta (DE3) and purification of recombinant His_6_-CMI1, His_6_-ICR1 and GST-ICR1 were carried out according to standard protocols using Ni-NTA (Qiagen) and Glutathione sepharose (GE) resins, as previously described (29, 57). His_8_-TEV-CMI1 was purified over Ni-NTA (Quiagen). Eluted fractions were passed through HiPrep 26/10 desalting column (GE Healthcare) with the extraction buffer (50mM sodium phosphate buffer pH 7.5, 300mM NaCl and 1mM DTT) to insure flushing of imidazole presence from the elution buffer (50mM sodium phosphate buffer pH 7.5, 300mM NaCl, 1mM DTT and 250mM imidazole). Eluted fractions were incubated overnight with His_6_-tagged TEV protease at 4°C followed by purification over a second Ni-NTA. The untagged CMI1 was collected from the flow through and concentrated with Amicon Ultra-15 with <u>m</u>olecular <u>w</u>eight <u>c</u>ut-<u>o</u>ff (MWCO) of 3kDa (Millpore) at 4,000 X *g* and 4°C to a final volume of ~500 μl. The concentrated protein samples were filtrated through Millex 0.22 μm syringe filter (MILLIPORE) and uploaded onto a gel filtration column of HiLoad 16/600 Superdex 200 pg (GE Healthcare) and eluted with a gel filtration column buffer (60 mM MOPS pH 7.2, 200 mM KCl and 2 mM DTT). Purified proteins were concentrated using Amicon Ultra-15 with MWCO of 3 KDa at 4,000 X *g* and 4°C, divided into aliquots, batch frozen in liquid nitrogen, and kept at −80°C until further use.

Anti-CMI1 antibodies were raised in rabbits. Ni-NTA purified His_6_-CMI was further purified by SDS-PAGE. The His_6_-CMI1 band was eluted from the gels and were used for rabbit immunization.

### *In vitro* ICR1-CMI1 and ICR1-ICR1 interaction assays

Pull-down of His_6_-CMI1 or HiS_6_-ICR1 with GST-ICR1: 1.2 μg GST-ICR1 or 0.4 μg GST were mixed with 100 μL of Phosphate Buffer Saline (PBS), 1% Triton X-100 and 10 μL of Glutathione sepharose slurry and incubated with shaking for 30 min at room temperature (RT). The beads were then washed 3X with PBS, 1% Triton X-100, and were adjusted in Ca^2+^/EGTA reaction buffer: 20 mM Tris-HCl pH 7.5, 5 mM CaCl_2_/10 mM EGTA, 0.1 mg/mL BSA, 200 mM NaCl, 1% Triton X-100. 0.5 μg His_6_-CMI1 were added for pull-down of His_6_-CMI1 by GST-ICR1. Alternatively, 0.09/0.45/1 μg His_6_-CMI1 and 0.5 μg His_6_-ICR1 were added for pull-down of HiS_6_-ICR1 by GST-ICR1. The reaction volumes were then adjusted to 100 μL with the respective buffer. The mixtures were incubated with shaking for 1 h at RT. Subsequently, the beads were precipitated and washed 1X with wash buffer 1: 20 mM Tris-HCl pH 7.5, 5 mM CaCl_2_/10mM EGTA, 0.1 mg/mL BSA, 1M NaCl, 1% Triton X-100 and 4X in wash buffer 2: 20 mM Tris-HCl pH 7.5, 5 mM CaCl_2_/10mM EGTA, 0.1 mg/mL BSA, 200 mM NaCl, 1% Triton X-100. The beads were then precipitated and resuspended in SDS-PAGE sample buffer and the proteins were resolved by SDS-PAGE (58).

Co-immuno precipitation of His_6_-ICR1 and His_6_-CMI1 with anti-CMI1 antibodies: His_6_-CMI1 and His_6_-ICR1, 1 μg of each, were incubated with shaking in 300 μL of Ca^2+^/EGTA reaction buffer for 1 h at RT. Subsequently, 1 μL of anti-CMI1 antibodies were added and the mixture was further incubated with shaking for 2 h at RT. 10 μL of Protein A beads (Adar Biotech #10165) slurry in Ca^2+^/EGTA reaction buffer were added and the mixture was further incubated with shaking for 1 h at RT. Subsequently, the beads were washed 3X with 1 mL ice cold Ca^2+^/EGTA reaction buffer, resuspended in SDS-PAGE sample buffer and proteins were resolved by SDS-PAGE. Proteins were detected by immunoblots decorated with mouse anti poly-His monoclonal antibodies (Sigma H-1029) and Goat anti mouse Horse Radish Peroxidase (HRP) conjugated secondary antibodies (BioRad).

### Circular dichroism (CD) spectroscopy

Protein samples were dialyzed overnight in buffers contained 10 mM Tris-H_2_SO_4_ pH 7.5, 25 mM KCl and 200 μM DTT. Buffers also contained CaCl_2_ and EDTA at different concentrations to obtain the desired Ca^2+^ free concentrations (Table 1). All protein samples and buffers were filtrated before use through Millex 0.22 μm syringe filter (MILLIPORE) or Stericup 0.22 μm vacuum filtration system (MILLIPORE), respectively. Protein concentration was determined using a Bradford assay standard curve for BSA. Cuvette path length was 0.1 mm and samples concentrations were 60 μM. Measurements were performed using a Chirascan CD spectrometer (Applied Photophysics), ranging between 180 nm to 260 nm at 21°C. Using the Pro-Data Viewer software (https://www.photophysics.com), each spectrum was averaged from five repeated scans. Then, raw data were corrected by subtracting the contribution of the buffer to the signal, subtracted data were smoothed (5 nm window) and exported to Excel. In Excel, data converted from observed ellipticity to <u>m</u>ean <u>r</u>esidue <u>e</u>llipticity (MRE) units using the following equation:

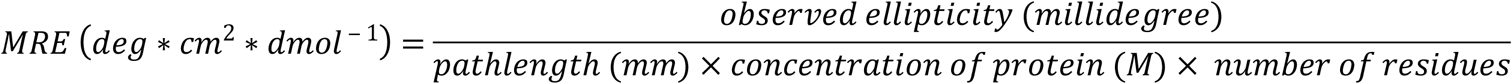

All measurements were repeated at least thrice. The α-helical content of sampled proteins was extracted from MRE values at 222 nm using the following equation (59):

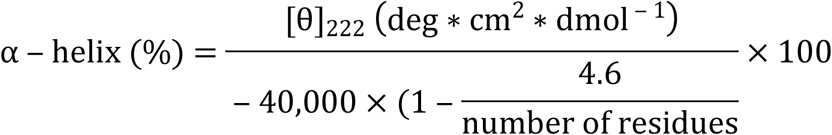

The α-helical content was averaged from the three repetitions and <u>s</u>tandard <u>e</u>rror (SE) was calculated as well.

**Table 1.** CaCl_2_ and EDTA composition in the CD spectroscopy buffers.

| Buffer name | 2 mM CaCl<sub>2</sub> | * 200 μM Ca<sup>2+</sup> free | * 20 μM Ca<sup>2+</sup> free | * 2 μM Ca<sup>2+</sup> free | * 200 nM Ca<sup>2+</sup> free | * 20 nM Ca<sup>2+</sup> free | * 2 nM Ca<sup>2+</sup> free | 1 mM EDTA |
| --- | --- | --- | --- | --- | --- | --- | --- | --- |
| CaCl<sub>2</sub> | 2 mM | 1199.9 μM | 1019.6 μM | 998.8 μM | 969.5 μM | 759.7 μM | 240.2 μM | - |
| EDTA | - | 1 mM | 1 mM | 1 mM | 1 mM | 1 mM | 1 mM | 1 mM |

\* Calculations were made using the WEBMAXC EXTENDED server (http://maxchelator.stanford.edu/webmaxc/webmaxcE.htm)

### Size Exclusion Chromatography coupled Multi-Angle Light scattering (SEC-MALS)

The SEC-MALS buffer containing 10 mM Tris-HCl pH 7.5, 25 mM KCl, 200 μM DTT and 2 mM CaCl_2_ was double filtrated through Stericup 0.22 μm vacuum filtration system (Millpore) and then through Whatman Anodisc 0.02 μm Filter Membranes (GE Healthcare). Protein samples were filtrated through Whatman Anotop 10 Plus 0.02 μm syringe filter (GE Healthcare) and their concentration was determined using a Bradford assay standard curve for BSA. Protein samples were injected into a Shodex KW404-4F column (Shodex) equilibrated overnight with the buffer above. The Agilent 1200 Series HPLC System (Agilent Technologies) was coupled with a DAWN HELEOS II light scattering detector (Wyatt Technology) and an Optilab rEX refractive index detector (Wyatt Technology). Molecular mass analyses were performed using the ASTRA software (https://www.wyatt.com/products/software/astra.html). Data were exported from the ASTRA software in order to build the graphs in Excel.

### Homology modeling of CMI1

Amino acids sequences of KIC and CMI1 were underwent a <u>m</u>ultiple <u>s</u>equence <u>a</u>lignment (MSA) using the MUSCLE algorithm (https://www.ebi.ac.uk/Tools/msa/muscle). The MSA results were converted into PIR format with the necessary adjustments to the solved crystal structure of KIC-KCBP complex (PDB ID code 3H4S) (36). A pairwise alignment of KIC with CMI1 was extracted from the MSA PIR format and run using the Modeller 9.19 (https://toolkit.tuebingen.mpg.de/#/tools/modeller). The built model of CMI1 was examined using the WHAT_CHECK (SAVES 5.0 server (http://servicesn.mbi.ucla.edu/SAVES)). The CMI1 model was visualized and edited using PyMOL (https://www.schrodinger.com/pymol) and Adobe Photoshop CS6.

### Yeast two-hybrid assays

*S. cerevisiae* strains Y190 and *PJ69-4a* were used as hosts. *pGAD-CMI1* or *pGAD-cmi1D85N* plasmids were co-transformed with *pGBT-ICR1/pGBT-ICR2/pGBT-ICR4* into yeast cells via a standard lithium acetate transformation protocol. Colonies expressing both plasmids were grown on a medium lacking leucine (Leu), tryptophan (Trp) and histidine (His) and containing 50 mM 3-Amino-1,2,4-triazole (3-AT). In addition, β-galactozidase activity assays were performed. Each test was carried out with at least four independent transformants. Assays with of site directed mutants in CMI1 and ICR1 and deletion mutants of ICR1 were carried out with *PJ69-4a* yeast. The <u>o</u>ptical <u>d</u>ensity in <u>600</u> nm (OD_600_) was measured and diluted into OD_600_ of 0.5. From this yeast suspension (referred as 1), four decimal dilution were made (1:10, 1:100, 1:10^3^ and 1:10^4^). From each dilution, a drop of 5 μl was placed on -LT (S2 Table) and - LTH (S3Table) with YNBx1, 2% glucose and 1mM <u>3</u>-<u>A</u>mino-1,2,4-<u>t</u>riazole (3AT) plate. The plates were incubated at 21°C.

### Plant protein co-localization assays

Co-localization assays were performed using transient expression of tested proteins by transforming *Agrobacterium tumefaciens GV3101 pMP90* cells harboring the respective plasmids into the abaxial side of *Nicotiana benthamiana* (*N. benthamiana*) leaf epidermis essentially as previously described (29, 60) with the following modifications. In cases were expression levels were too low for detection, *Agrobacterium* expressing the silencing suppressor protein p19 form tomato bushy stunt virus (61) were co-transformed added at a dilution OD_60_0 of 0.05. Following transformation plant were maintained in growth room for 48 hours prior to imaging.

### Immunostaining

Immunostaining of PIN1 and PIN2 in *Col-0* wild type *and cmi1* mutant roots was carried out essentially as previously described (62). Primary antibodies used in this study: anti-PIN1 (1:1000; sc-27163; Santa Cruz Biotechnology, Inc.) anti-PIN2 (1:400; N782248; NASC). Antirabbit Cy3 (1:600; CALTAG Laboratories, Invitrogen) and AlexaFluor 488 anti-rat (1:600; Invitrogen) were used as secondary antibodies. Fluorescence was observed using a Zeiss LSM780-NLO confocal microscope/multi photon microscope. Cy3 was observed by excitation at 543 nm and emission at 560 nm and AlexaFluor by excitation at 488 nm and emission at 499-519 nm emission. Quantification of PIN2 relocation was performed by scoring the number of cells with different PIN polarities.

### NaCl and RALF1 treatments and GUS staining

*cmi1 pCMI1::CMI-GUS* seedling were grown under long day conditions on 0.5X MS 1% sucrose lates. Then, seedlings were transferred to 0.5X MS 1% sucrose liquid medium and incubated for additional 3 hours. In turn, incubation media were removed and fresh 0.5X MS 1% sucrose mecia without (control) or supplemented with either 140 mM NaCl or 1 μM RALF1 peptide (Genscript) were added. For salt treatment seedlings were incubated for 3 and 6 hours. For treatments with RALF1 seedling were incubated for 4 hours and at 21°C followed by GUS staining for 3 hours at 37°C

### RNA Extraction and Quantitative PCR

For auxin induction experiments, seedlings were grown vertically on 0.5X Murashige Skoog (MS) supplemented with 1% sucrose for 5 days. Before treatment, seedlings were transferred to liquid 0.5X MS with 1% sucrose for 2 hours in a growth chamber. The 0.5X MS medium was then replaced with fresh 0.5X MS medium (mock) or 0.5X MS medium containing 1 μM of NAA. Following 2 hours incubation seedlings were frozen in liquid nitrogen and total RNA was extracted using the RNAeasy kit (Qiagen). RALF1 treatments, experiments were carried out essentially as described above but with treatments with 1 μM RALF1 for 0 (control) 15, 30, 45, 60, 90, 120 and 250 minutes. qPCR experiments were performed using the StepOnePlus™ Real-Time PCR System (Applied Biosystems). PP2A was used as a reference gene. The qPCR data was normalized to the reference gene. Three biological replicates with four technical replicates were carried out for each treatment. The qPCR program was as follows: 10 minutes at 95°C, followed by 40 cycles of 15 seconds denaturation at 95°C, 1 minute annealing, and elongation at 60°C. The results were analyzed using the StepOne™ software.

### Confocal Imaging

Confocal imaging was performed using Zeis780-NLO confocal laser scanning microscope (Zeiss, Jena, Germany) with 40X air, 20X, 40X and 63X water immersion objectives with NAs of 0.75, 0.8, 1.0 and 1.2, respectively. Protein tagged with eGFP or GFP were visualized by excitation with an argon laser at 488nm. Emission was detected with a spectral GaAsP detector set between 499nm to 552nm. Proteins tagged to mCherry or mRFP were visualized by excitation with an argon laser at 561nm and spectral GaAsP detector set between 579 nm to 632 nm. Image analysis was carried out with Zeiss ZEN 2012 (https://www.zeiss.com/microscopy/int/software-cameras.html) and Adobe Photoshop CS6 (https://www.adobe.com), Fiji (Image J) (https://fiji.sc/) and Imaris 8.4.1 (Bitplane).

### Microarray experiments

*Arabidopsis* seedlings were grown hydroponically for 6 days and subjected to auxin treatments as previously described (63). Roots were collected after 30 min of exposure to either 20 nM IAA (“low auxin”), 20 μM IAA (“high auxin”), or conditioned media (“mock”). Affymetrix ATH1 arrays were hybridized with probes generated from total RNA of four biologically independent samples per treatment. Data shown are mean signal values with standard deviation.

### *Arabidopsis* sample preparation and Ca^2+^ imaging

Experiments were carried out essentially as previously described (25). Surface-sterilized *Arabidopsis Ler* wild type or *cmi1* (3 independent lines for each) seeds expressing UBQ10-YC3.6 were plated on 0.5X strength MS medium (Duchefa) containing 1% (w/v) sucrose, solidified with 0.8% agar (Duchefa) (pH 5.8) and stratified for 2 d in the dark at 4°C. The plates were transferred to a growth chamber (16 h 22°C: 8 h 18°C, light: dark; 120-150 μmol m^−2^ s^−1^ light intensity) and seeds were grown vertically for 5-7 days. Single 5-7-days old *Arabidopsis* seedlings were placed inside a custom-made flow-through chamber (or perfusion chamber) containing imaging-buffer (5 mM KCl, 10mM MES and 10mM CaCl_2_, pH 5.8, adjusted with Tris). The seedling was fixed inside the chamber with cotton wool soaked in the imaging buffer as previously described (25, 64). The chamber was placed on the stage of an inverted ZEISS Axio observer (Carl Zeiss Microimaging GmbH, Goettingen, Germany) equipped with an emission filter wheel (LUDL Electronic Products, Hawthorne, NY, USA) and a Photometrics cool SNAPHQ2 CCD camera (Photometrics, Tucson, AZ, USA). A Zeiss Plan-APOCHROMAT 20/0.8 dry objective of the microscope was used for imaging. A xenon short-arc reflector lamp (Hamamatsu) with a 440-nm filter provided the excitation. Emission filters used were 485 nm (CFP) and 535 nm (YFP). A peristaltic pump was used for buffer circulation inside the flow through chamber with a flow rate of 1.5 ml min^−1^. YFP and CFP images were taken at 6-s intervals using the METAFLUOR software (Meta Imaging series 7.7; Molecular Devices, Downingtown, PA, USA). After monitoring the root in the buffer (continuous flow-through) for 2 min, the buffer was replaced by a buffer containing 10 μM NAA (Sigma Aldrich) for 7min.

### Ca^2+^ imaging data analysis

Offline calculation of the FRET ratio was performed using ImageJ64 software (http://rsb.info.nih.gov/ij/) with the RATIOPLUS plug-in. The intensities of CFP and YFP were measured from single CFP and YFP images as pixel intensity in arbitrary units. The ratio between YFP emission and CFP emission was calculated after background subtraction. We calculated the change in ratio R_t_-R_o_ or ΔR, where R_0_ is the basal ratio before application of the stimulus and R_t_ is the ratio at a specific time point. We normalized the ΔR to the basal ratio value (ΔR:R_0_) and plotted ratio graphs for each measurement. We aligned all the graphs to their first response point and plotted averaged ratio graphs.

### Analysis of Ca^2+^ responses

The Ca^2+^-peaks were divided into low, middle and high threshold peaks, depending on the ratio change presented as height of the peak from the base. Peaks with a ratio change of 0 to 0.1 were considered as low threshold peaks, a ratio change of 0.1 to 0.15 as intermediate and the peaks with ratio changes higher than 0.15 were considered as high threshold peaks. Percentage was then calculated. 15 to 17 seedlings were analyzed for each genotype. The average ratio graphs were calculated from six to seven measurements.

### High-resolution Ca^2+^ imaging

High-resolution imaging was performed as previously described (64), with a Leica DMI 6000B inverted microscope equipped with a Leica TCS SP5 laser scanning device and HDy, using the Leica confocal software (Leica Application Suite - Advanced Fluorescence 2.6.0.7266; Leica Microsystems, Wetzlar, Germany). For excitation, an argon laser with a 458 nm line was used. The CFP and fluorescence energy resonance transfer (FRET) emissions were collected at 473–505 and 526–536 nm, respectively. Images were acquired with a 25x objective (HCW RAPO L 25.0 x 0.95 water). Image acquisition was conducted as follows: scanning speed (400 Hz), image dimension (512 × 512), pinhole (2-4 airy unit) and line average (4). YFP and CFP images were acquired as a time series in a 6 s interval. Offline calculation of the FRET ratio was performed using ImageJ RATIOPLUS plug-in.

### Data analysis and statistics

The measurements of roots, hypocotyls and root hairs or fluorescence intensity were performed using Image J. The means and the standard errors (SE) were calculated using Excel; the significance (p_values_) was calculated using SPSS.

## Acknowledgements

We thank the Manna Center for Plant Biology at Tel Aviv University for support and Jacqueline Wyatt and Sigal Lazar for editing. This research was supported by the Israel Science Foundation (grant nos. ISF 827/15, ISF-UGA 2739/16), The Israel Center for Research Excellence on Plant Adaptation to Changing Environment (I-CORE 757-12) to SY, The German Research Foundation Germany-Palestinian Authority-Israel Trilateral Program (DFG KU 931/13-1) to JK and SY and Howard Hughes Medical Institute, the Gordon and Betty Moore Foundation and the NIH (GM43644) to ME.

## Supplemental Figures and Tables Captions

### S1 Fig. Amino acid sequence of CMI1, its induction by auxin and its Ca^2+^ dependent pull down by ICR1

(A) The amino acid sequence of CMI1. The loop region of the single EF-hand is underlined and the D85 critical for Ca^2+^ binding is highlighted in red. (B) Protein immuno blot decorated with anti polyHis antibodies showing that pull-down of His_6_-CMI1 by GST-ICR1 is specific and Ca^2+^-dependent. (C) Coomassie brilliant blue stained SDS-poly acrylamide gel showing specified *E. coli* expressed and purified recombinant proteins used for the pull-down and immuno precipitation assays (Figure 1). Numbers denote M_*r*_ in kDa.

### S2 Fig. CMI1 exist as a monomer in solution

A SEC-MALS elution profile of 2 μg CMI1 in 2 mM Ca^2+^ solution. CMI1 eluted as a single peak with a molecular mass (red line) corresponding to a monomeric form. The profile is identical to that obtained with 4 μg protein (Fig. 2E)

### S3 Fig. CMI1 displays weak self-interaction in yeast two-hybrid assays

Yeast two-hybrid assays were carried out in the LexA system. In CMI1 self-interaction assays, weak XGal activity was evident after 48 hours. Strong XGal activity was observed in assays with ICR1 and no activity with the vector control.

### S4 Fig. Interaction between CMI1 and icr1W266Q is similar to icrW266A

-LTH: Leu, Trp and His dropout medium. -LT: Leu and Trp dropout medium. Numbers above the panel denote dilution order.

### S5 Fig. ICR1 is localized on MTs

(A-C) ICR1-mCherry (ICR1) is colocalized to MTs with Tubulin6-GFP (TUA6) MTs marker on MTs. (D-F) Localization of ICR1 and GFP-CMI1 (CMI1) on MTs is sensitive to the anti MTs drug oryzalin. O/L mCherry/GFP overlay. Bar: 20μm

### S6 Fig. CMI1 is localized in the plasma membrane, cytoplasm and nuclei and sensitivity of its expression to salt stress

(A) Subcellular localization of mRFP-CMI1 in *Arabidopsis* cotyledon pavement cells. C-cytoplasm. N-nuclei, M-plasma membrane. Localization of mRFP-CMI1 in the plasma membrane can be seen following plasmolysis (right panel). (B) Protein immuno blot decorated with anti-CMI1 antibodies showing the distribution of CMI1 between the soluble and insoluble fraction in the specified tissue samples. (C) The sensitivity of the anti-CMI1 antibodies as determined by protein immuno blot of the specified amounts of His_6_-CMI1.

### S7 Fig. Expression of CMI1 in the root tip and its induction by auxin

(A) The root tip of *pCMI1>>mRFP-CMI1*. (B) qPCR showing induction of CMI1 expression 6 h after treatment with mock or 10 μM IAA.

### S8 Fig. CMI1-GUS can complement root growth inhibition in *cmi1* knockout plants

(A) Primary root length of 7 days-old seedlings. Error bars are SE. Representative hypocotyls (B) and root hairs (C) used for quantifications presented in Figure 5L and K, respectively.

### S9 Fig. Lateral root development in wild type and *cmi1*

(A) Wild type Lateral root initials (LRIs) at different developmental stages. (B) *cmi1* LRIs. Note the abnormal LRI patterning. The developmental stages of the LRIs are noted.

### S10 Fig. PIN2 auxin efflux transporter is altered in *cmi1*

(A) Immunolocalization of PIN1 in the endodermis (en) and PIN2 in the cortex (co) and the epidermis (ep) in *Col-0* (WT) and *cmi1*. Arrowheads highlight the basal localization of PIN2 in wild type cortex and apical and apolar localization in *cmi1* cortex. (B) Quantitative analysis of PIN2 distribution. Scale bar 20 μm. Error bars SE.

### S11 Fig. Interaction assays of CMI1 with auxin transport and signaling proteins in yeast 2-hybrid assays

(A) CMI2/PBP1 but not CMI1 interact with PID. (B and C) CMI1 was used as a bait in with prey as labeled in the tables. (D) Assays as labeled in the table. Note that the positive interaction between CMI1 and ARF5 (B) likely resulted from ARF5 self-activation (D).

### 12 Fig. CMI1 expression is repressed by NaCl and rapidly and transiently induced by RALF1

*cmi1 pCMI1::CMI1-GUS* seedling were treated with mock (control) or 140 NaCl solution for indicated times (A) or for 4 hours with mock (control) or 1 μM RALF1 (B) and (C). (D) qPCR of CMI1 following treatments with 1 μM RALF1 for indicated time points.

**S1 Table. Plasmids used in this study**

**S2 Table. Materials used in this work**

**S3 Table. *Arabidopsis thaliana* lines used in this study**

